# An essential host dietary fatty acid stimulates TcpH inhibition of TcpP proteolysis enabling virulence gene expression in *Vibrio cholerae*

**DOI:** 10.1101/2022.04.28.489952

**Authors:** Lucas M. Demey, Ritam Sinha, Victor J. DiRita

**Author notes:** Address corresponding to Victor J. DiRita, **Email:**. **Author Contributions:** Conceptualization: Lucas Demey and Victor DiRita; Methodology: Lucas Demey; Investigation: Lucas Demey and Ritam Shina; Writing – Original Draft: Lucas Demey; Writing – Review & Editing: Lucas Demey, Ritam Shina, and Victor DiRita; Visualization: Lucas Demey; Project Administration: Victor DiRita; Funding Acquisition: Victor DiRita.

## Abstract

*Vibrio cholerae* is a Gram-negative gastrointestinal pathogen responsible for the diarrheal disease cholera. Expression of key virulence factors, cholera toxin and toxin-coregulated pilus, is regulated indirectly by two single-pass membrane-localized transcription regulators (MLTR), ToxR and TcpP, that promote expression of the transcription activator *toxT*. TcpP abundance and activity are controlled by TcpH, a single-pass transmembrane protein, which protects TcpP from a two-step proteolytic process known as regulated intramembrane proteolysis (RIP). The mechanism of TcpH mediated protection of TcpP represents a major gap in our understanding of *V. cholerae* pathogenesis. Absence of *tcpH* leads to unimpeded degradation of TcpP *in vitro* and a colonization defect in a neonate mouse model of *V. cholerae* colonization. Here, we show that TcpH protects TcpP from RIP via direct interaction. We also demonstrate that a dietary fatty acid, α-linolenic acid, promotes TcpH-dependent inhibition of RIP via co-association of TcpP and TcpH molecules within detergent-resistant membranes (DRMs; also known as lipid rafts) in a mechanism requiring the TcpH transmembrane domain. Taken together our data support a model where *V. cholerae* cells use exogenous α-linolenic acid to remodel the phospholipid bilayer *in vivo*, leading to co-association of TcpP and TcpH within DRMs where RIP of TcpP is inhibited by TcpH, thereby promoting *V. cholerae* pathogenicity.

**Significance Statement:** *V. cholerae* continues to pose a significant global burden on health infection millions of people every year resulting in ∼100,000 deaths annually. The importance of *toxT* gene expression in *V. cholerae* pathogenesis has been well established. Our results show that TcpP, one of the major regulators of *toxT* gene expression, is protected from proteolysis by TcpH, via direct interaction, in the presence of α-linolenic acid, an essential dietary fatty acid. Here we identify a physiological relevant host factor that stimulates *V. cholerae* pathogenicity via TcpH-dependent antagonism of TcpP proteolysis.

## Introduction

*V. cholerae* tightly regulates expression of its virulence factors, such as cholera toxin (CtxAB) and the toxin co-regulated pilus (TcpA-F) to reach the optimal site of infection, the crypt of intestinal villi (1–6). Transcription of these essential virulence factors is regulated by ToxT, an AraC-like transcription factor (7–10). Similarly, transcription of *toxT* is highly regulated and positively stimulated by TcpP and ToxR, two membrane-localized transcription regulators (MLTRs) (11–14).

TcpP and ToxR are bitopic membrane proteins that each contain a cytoplasmic DNA-binding domain, a single transmembrane domain, and a periplasmic domain. Both ToxR and TcpP directly bind to the promoter region of *toxT*, at -180 to -60 and -55 to -37, respectively (9, 15, 16). While ToxR directly binds to the *toxT* promoter, ToxR alone is unable to directly stimulate *toxT* expression (9). However, TcpP is required for *toxT* expression, presumably because TcpP facilitates transcription through direct interaction with RNA polymerase due to its binding sequence being near the -35 site (9, 15). Unlike ToxR, transcription of *tcpP* is tightly regulated by multiple transcription factors, further demonstrating the critical importance of TcpP (17–24).

TcpP is also post-translationally regulated by two proteases, Tail-specific protease (Tsp) and YaeL, through a process known as Regulated Intramembrane Proteolysis (RIP) (25–27). RIP is a form of gene regulation conserved across all domains of life that allows organisms to rapidly respond to extracellular cues, commonly by liberating a transcription factor or a sigma factor, from membrane sequestration (28). Two well-characterized systems controlled by RIP mechanisms are the extracytoplasmic stress response in *E. coli* and sporulation in *Bacillus subtilis*. These systems require RIP of RseA and SpoIVFB, respectively, to release their respective sigma factors (σ^E^ and pro-σ^K^) from the membrane and activate gene expression (29–35). Similarly, both systems have their respective TcpH analog, RseB and BofA, which function to prevent RIP of RseA and SpoIVFB via different mechanisms (30, 36–41). Regulation of TcpP by this mechanism diverges from these systems because transcription activity of TcpP is not activated by RIP but is rather inactivated by RIP, removing TcpP from the cytoplasmic membrane and thereby decreasing *toxT* transcription (25–27).

Our current understanding of RIP of TcpP is limited. Under RIP-permissive conditions *in vitro* (e.g. LB pH 8.5, 37°C), TcpP is sensitive to proteolysis by tail-specific protease (Tsp; site-1 protease), and subsequently by YaeL protease (site-2 protease) (25–27). RIP of TcpP is inhibited by its associated protein, TcpH, under specific *in vitro* conditions (e.g. LB pH 6.5, 30° C) (25–27). In cells lacking TcpH, TcpP is constitutively sensitive to RIP (25–27). However, the mechanism by which TcpH inhibits RIP and how TcpH-dependent RIP inhibition is modulated by extracellular stimuli remains unknown.

Detergent-resistant and detergent-soluble membranes (DRM and DSM, respectively) (i.e., lipid-ordered and lipid-disordered membrane domains) are known to form in both eukaryotic and prokaryotic organisms (42–48). In prokaryotes, lipid-ordered membrane domains are small phospholipid domains that exist within both inner and outer membranes (43, 44, 48). They are composed of saturated phospholipids and hopanoids (or cholesterol in eukaryotic cells) that tightly interact, resulting in a structured membrane region with low fluidity. Conversely, lipid-disordered membrane domains are enriched in unsaturated phospholipids resulting in high fluidity (42–44, 46– 55). Due to these differences lipid-ordered and lipid-disordered membrane domains can be separated based on solubility in non-ionic detergents, and we refer to them as detergent-resistant membranes (DRM) and detergent-soluble membranes (DSM), respectively.

In this report we provide evidence that TcpH protects TcpP from RIP via direct interaction. Furthermore, we explore the role of the membrane, specifically lipid-ordered and disordered domains, in regulating TcpP-TcpH association. Our data suggest that *in vivo* TcpP and TcpH preferentially associate with DRMs. This leads to enhanced inhibition of RIP by TcpH, thereby resulting in elevated TcpP levels, and *toxT* transcription. We also show that utilization of exogenous α-linolenic acid, a long chain poly-unsaturated fatty acid present *in vivo*, stimulates TcpP and TcpH association within DRMs. Data generated here support a model where, once *V. cholerae* cells enter the gastrointestinal tract, cellular uptake of α-linolenic acid results in modification of the phospholipid profile and leads to an increase the abundance of TcpP and TcpH molecules within DRMs thereby stimulating inhibition of RIP.

## Results

### Altering the transmembrane and periplasmic domains does not disrupt TcpH *in vitro* activity

To identify regions within TcpH critical for its role in protecting TcpP from RIP we constructed chimeric transmembrane domain fusions (TM) and periplasmic TcpH deletion constructs (Peri). Two TM and one Peri constructs [_ToxS_TcpH, _EpsM_TcpH, and TcpH_Δ103-119_, respectively] were constructed, and the allele encoding each was recombined into the *V. cholerae* genome so as not disrupt the *tcpP* coding sequence, and under normal *tcpPH* transcriptional control (Figure 1A). Growth of the resulting strains was unaffected in comparison with wild-type *V. cholerae* in virulence inducing (Vir Ind) conditions (Figure S1A). We evaluated the constructs also by measuring TcpP levels, *toxT* transcription, and TcpA and CtxB production in vitro (Figure 1B-D and Figure S1B). All TcpH constructs protected TcpP, supported *toxT* transcription, and regulated virulence factor production similar to WT TcpH and better than Δ*tcpH* (Figure 1B-D).

**Figure 1:**
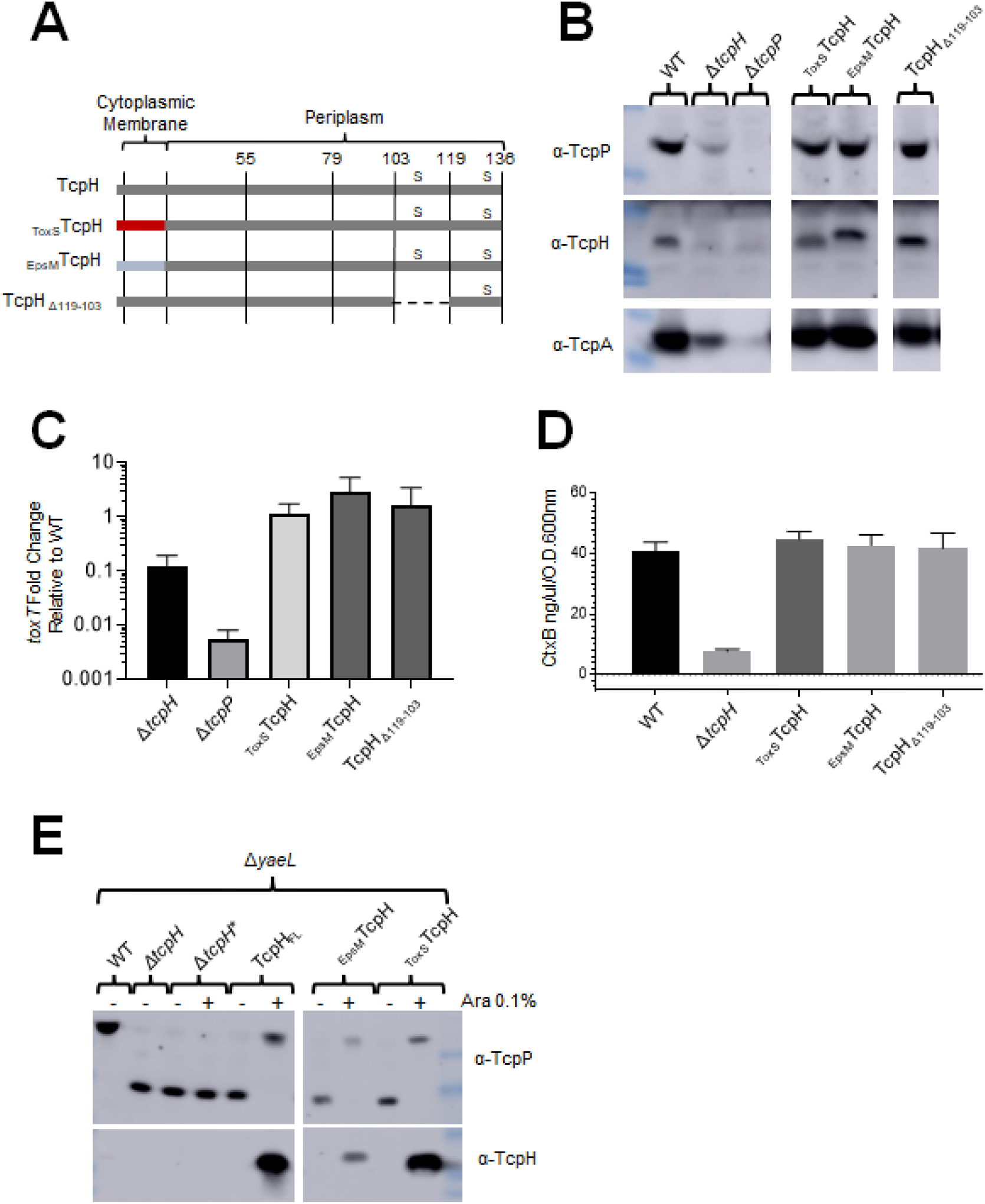
TcpH transmembrane and periplasmic constructs protect TcpP, support *toxT* expression, and virulence factor production. A) Diagram of TcpH transmembrane constructs (_EpsM_TcpH and _ToxS_TcpH) and periplasmic construct (TcpH_Δ119-103_). TcpH has a single transmembrane domain (also a Sec signal sequence), at its N-terminus, and two periplasmic cysteine residues (C114 and C132), represented by “s”. The transmembrane domain of TcpH was replaced with the transmembrane domain of ToxS (_ToxS_TcpH) and EpsM (_EpsM_TcpH) as both ToxS and EpsM are known to be localized to the cytoplasmic membrane with similar domain topology as TcpH (139, 140). In-frame deletion of periplasmic residues are indicated by a dashed line. B-D) *in vitro* characterization of TcpH transmembrane and periplasmic chromosomal constructs grown under virulence inducing conditions. B) Western blots of whole-cell lysates probed with α-TcpP (top), α-TcpH (middle), and α-TcpA (bottom). C) Average *toxT* transcription of three biological replicates, determined via Δ ΔCT method. *toxT* fold change is relative to WT *V. cholerae*. B) CtxB levels, measured via ELISA, in culture supernatants collected from cultures incubated with *V. cholerae* cells cultured in virulence inducing conditions for 24hrs. Error bars represent standard error of the mean. See Figure S1B for the unmodified western blots in panel B. E) Western blots of spheroplast fractions (cytoplasm and cytoplasmic membrane fractions). TcpH transmembrane constructs (_ToxS_TcpH and _EpsM_TcpH) and native TcpH were expressed from pBAD18 in Δ*tcpH* Δ*yaeL* background under virulence inducing conditions for 6hrs. All strains, excluding WT, are Δ*tcpH* Δ*yaeL*. Δ*tcpH*\* harbors pBAD18 (empty vector).

While the TM TcpH complement a Δ*tcpH* mutant by supporting higher levels of TcpP *in vitro*, we sought to determine if the TM TcpH constructs specifically inhibited RIP of TcpP. In the absence of TcpH, TcpP is sensitive to degradation and undergoes RIP. Loss of both *tcpH* and *yaeL* leads to accumulation of TcpP*, an (functional) intermediate degradation product formed by cleavage of TcpP by Tsp alone and which serves as the substrate for YaeL (56). TcpP* lacks most of its periplasmic domain and therefore has a lower molecular weight (∼17 KDa) compared to TcpP (∼29 KDa), thus enabling us to determine the RIP status of TcpP via western blot. When TcpH is active and RIP is thereby inhibited, we observe full-length TcpP and no TcpP*. When TcpH, _ToxS_TcpH, or _EpsM_TcpH constructs were ectopically expressed in a *ΔtcpH/ ΔyaeL* we observed only full-length TcpP and no TcpP* (Figure 1E). Taken together, our data indicate that replacing the transmembrane domain does not disrupt TcpH function *in vitro*.

### TcpH TM domain is critical for colonization of infant mice

The TM and Peri domain of TcpH can withstand considerable modifications and maintain function in *V. cholerae* as determined by *in vitro* experiments. To test whether this was the case *in vivo*, we infected infant mice with strains expressing TcpH TM and Peri constructs (Figure 2A). Despite their wild-type activity *in vitro*, strains expressing _ToxS_TcpH, and _EpsM_TcpH colonized infant mice to levels significantly lower than did wild type, more closely resembling those of the Δ*tcpH* strain (Figure 2A). TcpH_Δ103-119_ supported the same level of TcpH-dependent virulence gene expression *in vitro* as both _ToxS_TcpH and _EpsM_TcpH, but colonized infant mice to a similar degree as wild type (Figure 2A). The inocula of _ToxS_TcpH and _EpsM_TcpH used to infect infant mice produced similar levels of TcpA compared to wild type (Figure S2A).

**Figure 2:**
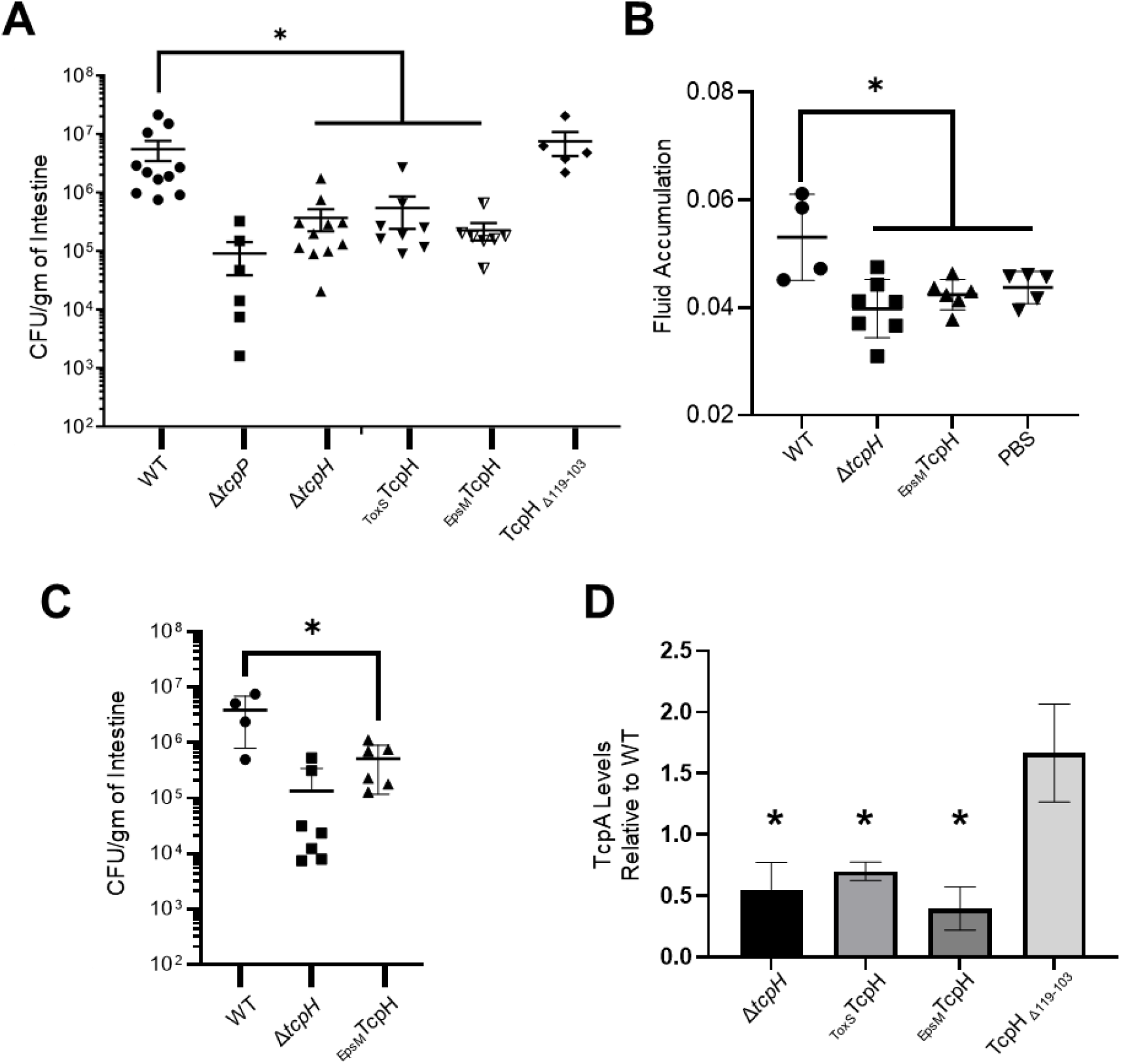
TcpH transmembrane domain is critical *in vivo*. A) Colony forming units (CFU’s) per gram of 3-6 day old infant mouse intestine 21hrs post infection (1×10^6 inoculum dose). The horizontal line indicates the average CFU/gm of intestine and is an average of 5-11biological replicates. B and C) Fluid accumulation and CFU’s per gram of infant mouse intestine 18hrs after infection (1×10^8 inoculum dose). D) Relative TcpA levels after 21hrs of aerobic growth in sterile adult mice fecal media (9% w/v). TcpA levels were determined via densitometry, calculated using ImageJ. Averages represent three or more biological replicates. Error bars represent standard error of the mean in panels A-C and standard deviation of the mean in panel D. A mann-whitney U test was used to determine statistical significance in panel A. A two-tailed Student’s t-test was used to determine statistical significance in panel B and C. A one-tailed Student’s t-test was used to determine statistical significance in panel D. * Indicates a p-value less than 0.05.

We hypothesized that the colonization defects of the TM TcpH constructs were likely due to an inability of strains lacking the natural TcpH transmembrane domain to express colonization factors particularly TcpA – *in vivo*. To test this we measured disease signs, such as fluid accumulation, which requires a higher infectious dose compared to colonization experiments (e.g., 1*10^8 vs. 1*10^6). Mice infected with _EpsM_TcpH exhibited lower fluid accumulation compared to those infected with wild type *V. cholerae* and, despite a higher infectious dose, _EpsM_TcpH still was unable to colonize infant mice to wild type levels (Figure 2BC). These data support our hypothesis that _EpsM_TcpH is unable to colonize infant mice due to an inability to support virulence factor production both TcpA and CtxAB – *in vivo*.

To determine whether the presence of other microbes in the gastrointestinal tract might influence the ability of strains expressing TcpH with altered TM domains to support virulence gene expression, we cultured wild type and the TcpH constructs (TM and Peri) aerobically in both filter-sterilized and non-filtered mouse fecal media for 21 hrs at 37°C (Figure S2BC). All strains exhibited similar growth rates and final cell densities in both filtered and non-filtered fecal media (Figure S2BC). We quantified TcpA levels in cell lysates after 21 hours of growth in sterile mouse fecal media. While the growth rates were very similar between wild type and strains expressing altered TcpH proteins, the strains expressing _ToxS_TcpH and _EpsM_TcpH produced TcpA levels below that of wild type (Figure 2D). The strain expressing the TcpH protein with a periplasmic deletion was unaffected for TcpA expression (Figure 2D). Taken together, these data suggest that the TcpH transmembrane domain is critical for TcpH in the gastrointestinal tract to protect TcpP from RIP, thereby supporting downstream virulence factor production. Due to their wild type levels of colonization and ability to support wild type levels of TcpA synthesis in mouse fecal media we excluded the TcpH Peri construct from further experiments.

### *toxT* transcription is enhanced with crude bile and is dependent on the TcpH transmembrane domain

Data presented here and elsewhere indicate that TcpH-dependent RIP inhibition is affected by different *in vitro* and *in vivo* environmental signals and that the transmembrane domain of TcpH is critical for that function (25–27). *Vibrio* species can use exogenous fatty acids present in bile via the VolA and FadL/FadD pathways (57–61), resulting in modification of membrane phospholipid composition (61, 62). Phospholipid remodeling in *V. cholerae* can influence growth rate, biofilm formation, and motility (61, 62). Given that TcpH and TcpP require membrane localization, we sought to determine whether phospholipid changes, perhaps stimulated by fatty acids present in the gastrointestinal tract, would influence TcpH-dependent inhibition of TcpP RIP.

We measured *toxT* expression from a transcription reporter (pBH6119-*toxT::GFP*) in cells grown in media supplemented with Bovine Crude Bile (0.4%), which contains various fatty acids that can be incorporated into the bacterial membrane (61), Transcription of *toxT* from this reporter was elevated in the presence of crude bile in cells expressing wild type TcpH, but not in cells expressing TcpH constructs with transmembrane domains from EpsM or ToxS (Figure S3A). This suggested that native TcpH responds to changes in phospholipid composition to inhibit RIP, and that TcpH with the altered transmembrane domain is unable to sense and/or respond to the same change. As a negative control, we measured *toxT* transcription under non-inducing conditions known to stimulate RIP of TcpP (25–27), and in these conditions *toxT* expression was indeed reduced (Figure S3A). In addition, we measured *toxT* expression in Δ*tcpP* and Δ*tcpH* cells with and without crude bile present, observing no increase in *toxT* expression (Figure S3A). This confirms that the conditions used here do not simply promote TcpP function in the absence of TcpH.

We also measured *toxT* transcript levels directly via RT-qPCR with RNA isolated from wild type cells grown in the presence of crude bile (Figure S3B), and observed a similar increase in *toxT* transcription. Lastly, we found that cells expressing native TcpH or TcpH with altered TM domains grew with similar rates in crude bile-supplemented media used for these experiments (Figure S1C). Taken together, these data support a model that TcpH responds to host stimuli, specifically fatty acids or constituents of crude bile, through a mechanism requiring its native TM, and antagonizes RIP of TcpP, leading to increased *toxT* transcription.

### α-Linolenic acid enhances *toxT* expression by promoting TcpH dependent enhanced RIP inhibition

Crude bile is a mixture of saturated and unsaturated fatty acids, as well as bile salts (e.g., cholate and deoxycholate). To determine whether bile salts or fatty acids in crude bile were responsible for elevated *toxT* transcription in WT, we supplemented virulence-inducing media with cholate/deoxycholate (Purified Bile) (100µM of each), palmitic acid (500µM), stearic acid (500µM), linoleic (500µM), α-linolenic acid (500µM), arachidonic acid (500µM), and docosahexaenoic acid (500µM). Using the *toxT*::GFP transcription reporter plasmid, we observed elevated *toxT* transcription in wild type cells with only crude bile or α-linolenic acid present (Figure 3A and Figure S3A). Increased *toxT* expression with crude bile or α-linolenic acid was not observed in *ΔtcpH* or *ΔtcpP* cells (Figure S3A), demonstrating that TcpH is still needed to inhibit RIP and TcpP is necessary to promote *toxT* transcription in the presence of these compounds. None of the purified components of crude bile resulted in statistically significant increased levels of *toxT* expression in cells expressing _EpsM_TcpH or _ToxS_TcpH (Figure S3A). Expression of *toxT* responds in dose-dependent fashion to the presence of α-linolenic acid (Figure S3C).

**Figure 3:**
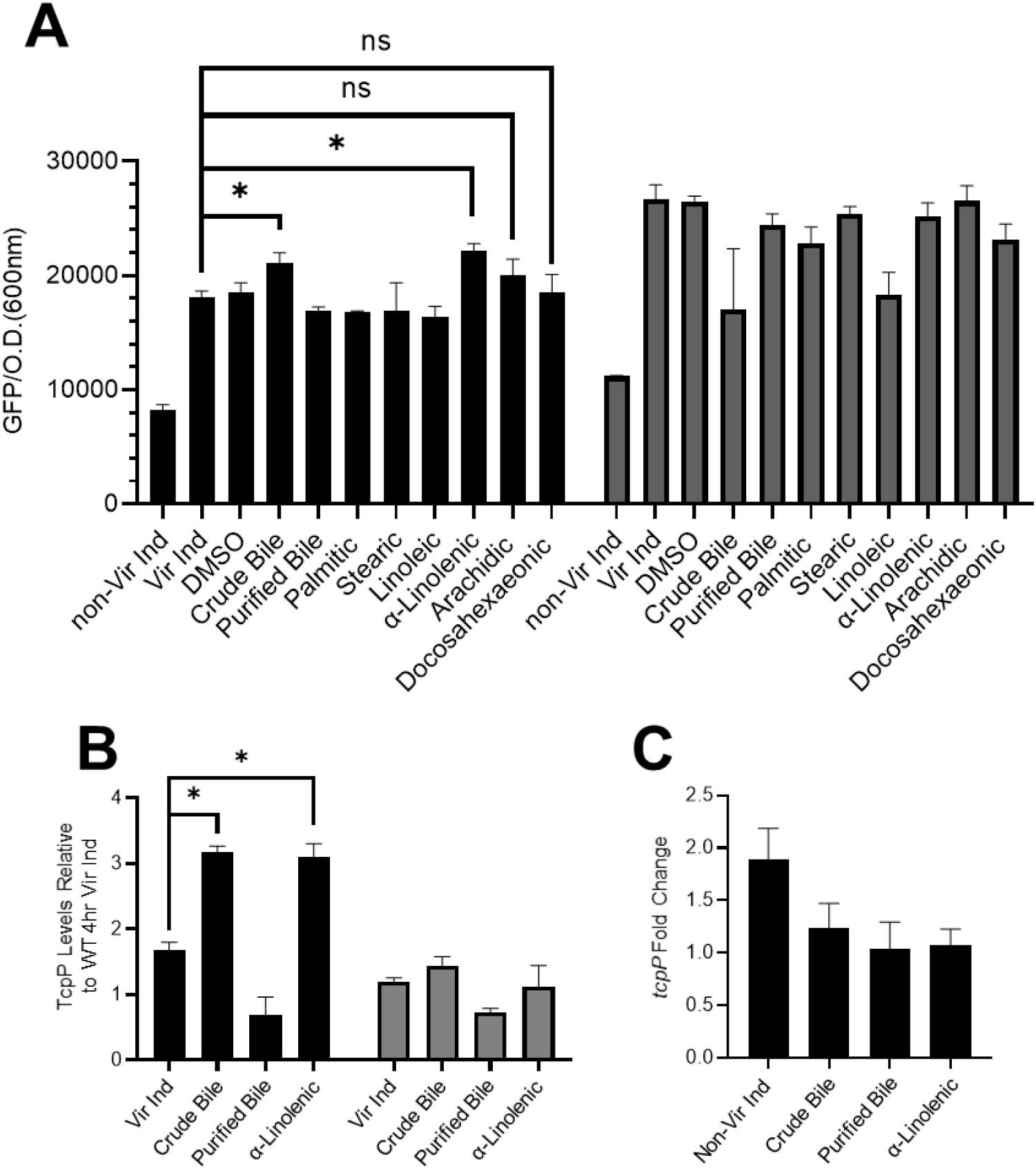
α-Linolenic acid stimulates a TcpH transmembrane dependent increase in *toxT* transcription, elevated TcpP levels, and does not increase *tcpP* expression. A) *toxT* expression in WT (black bars) and _EpsM_TcpH (gray bars) was determined using a plasmid based *toxT::GFP* transcription reporter. *toxT* transcription was determined by measuring GFP fluorescence (excitation 488 nm and emission 515 nm) and optical density (600 nm). The data here are an average of three or more biological replicates, and error bars represent the standard error of the mean. B) TcpP levels in WT (black bars) and _EpsM_TcpH (gray bars) relative to WT cells cultured under virulence inducing conditions. Densitometry, calculated by ImageJ, was used to determine relative abundance of TcpP. Error bars represent standard error of the mean. A and B) Two-tailed Student’s t-test was used to determine statistical significance. * indicates a p-value of less than 0.05. C) *tcpP* transcription in WT *V. cholerae* cells using RT-qPCR, determined via Δ ΔC_T_ method. *tcpP* transcription is relative to WT Vir Ind. Error bars represent standard error of the mean.

In addition to the results obtained with the *toxT::GFP* reporter, we measured *toxT* mRNA levels using RT-PCR in WT cells grown under the same conditions. Consistent with the reporter plasmid data, we observed elevated *toxT* mRNA in the presence of α-linolenic acid (∼2.5 fold) (Figure S3B). There was no difference in growth rate in cells expressing either native TcpH or the TcpH TM constructs when cultured with purified bile or α-linolenic acid (Figure S1DE).

We reasoned that enhanced *toxT* transcription in the presence of crude bile or α-linolenic acid was due to enhanced inhibition of RIP, leading in turn to elevated levels of TcpP. Thus, we quantified TcpP levels under virulence inducing conditions supplemented with crude bile or α-linolenic acid (Figure 3B). TcpP levels in wild type cells were significantly elevated in the presence of crude bile or α-linolenic acid (Figure 3B). In contrast, growth in α-linolenic acid had no effect on TcpP levels in cells expressing _EpsM_TcpH grown (Figure 3B). Loss of TcpH led to degradation of TcpP under all conditions indicating that Tsp and YaeL activity is not inhibited by the addition of crude bile or α-linolenic acid (data not shown). We conclude that i) elevated *toxT* transcription in the presence of crude bile or α-linolenic acid is due to enhanced inhibition of RIP via TcpH and ii) altering the phospholipid composition of the cells with exogenous crude bile or α-linolenic acid enhances TcpH function in RIP inhibition through a mechanism that requires the native transmembrane domain.

As TcpP levels are elevated upon supplementation of crude bile or α-linolenic acid, we considered it possible that elevated *tcpP* transcription could contribute to elevated TcpP levels. One possible mechanism is that *tcpP* transcription is directly influenced by α-linolenic within the cytoplasm. Prior studies have shown that linoleic acid can rapidly diffuse into the cytoplasm of *V. cholerae* where we reasoned it might influence *tcpP* gene expression (63, 64). To determine if *tcpP* transcription is influenced by crude bile or α-linolenic acid we measured *tcpP* transcription in wild type *V. cholerae* cells using both RT-qPCR and a transcription reporter, *tcpP::lacZ*. Neither crude bile nor α-linolenic acid supplementation led to increased *tcpP* transcription (Figure 3C and Figure S4A). These data indicate that crude bile and α-linolenic acid influence TcpP levels post-transcriptionally supporting the hypothesis that these conditions lead to elevated TcpP by enhanced TcpH-dependent RIP inhibition.

*V. cholerae* can use exogenous fatty acids, including α-linolenic acid, for *de novo* phospholipid biosynthesis, thereby modifying its phospholipid composition (61, 62). To determine whether *V. cholerae* cells incorporate α-linolenic acid into the cytoplasmic membrane as phospholipids under our growth conditions, we analyzed the fatty acid profile of phospholipids from *V. cholerae* cells cultured with and without α-linolenic acid (Figure S4B). In the presence of α-linolenic acid more than 80% of acyl chains within *V. cholerae* were 18:3. This is consistent with prior published data (61, 62) and demonstrates that under our conditions *V. cholerae* cells are remodeling the fatty acid content of their phospholipids. Given that the vast majority of fatty acids detected are 18:3 and prior studies indicate that *V. cholerae* does not synthesize 18:3 fatty acids under standard laboratory conditions (65, 66), these data suggest that *V. cholerae* cells are utilizing exogenous α-linolenic acid for phospholipid synthesis (Figure S4B).

We next assessed whether cells expressing native TcpH or _EpsM_TcpH were equally capable of using exogenous fatty acids. This is to rule out any unforeseen, and unlikely, issue with fatty acid metabolism in cells expressing non-native TcpH that might mislead our conclusions about the importance of the native TcpH transmembrane domain in regulating TcpH function. We cultured cells with cerulenin, an inhibitor of *de novo* fatty acid synthesis, or with cerulenin plus exogenous fatty acids (67–70). Cerulenin alone led to a growth defect of cells irrespective of which form of TcpH was being expressed (Figure S4C). Inclusion of unsaturated fatty acids restored partial growth of both (Figure S4C). These data indicate that incorporation of fatty acids into the phospholipid bilayer is unaltered in cells expressing _EpsM_TcpH, supporting our conclusion that the defect of this protein in protecting against RIP is due to its altered transmembrane domain.

### Co-Association of TcpP and TcpH with detergent resistant membranes is required for enhanced RIP inhibition

Our work demonstrates that under conditions that modify phospholipid composition, TcpP levels are enhanced, and *toxT* transcription is increased. Elevated levels of TcpP are likely due to enhanced inhibition of RIP by TcpH rather than increased *tcpP* transcription, and this inhibitory function requires the native TcpH TM domain. In addition to α-linolenic acid, arachidonic and docosahexaenoic acid modify phospholipid composition in *V. cholerae* (61). Despite causing similar changes to the phospholipid profile, these polyunsaturated fatty acids do not have a significant effect on *toxT* transcription (Figure 3A and Figure S3A). These data indicate the phospholipid profile does not predict TcpH-dependent inhibition of RIP. Exogenous fatty acids can be utilized directly as acyl chains in *de novo* phospholipid synthesis (71, 72). Thus, while gross phospholipid composition can remain similar with supplementation of α-linolenic, arachidonic, and docosahexaenoic acid, (i.e., relative abundance of cardiolipin, phosphatidylglycerol, and phosphatidylethanolamine) the overall biophysical properties of the cytoplasmic membrane (i.e., membrane fluidity) can differ due to differences in acyl chain composition. We reasoned that the differences in observed TcpH-dependent enhanced RIP inhibition could be due to differences in the biophysical properties of the cytoplasmic membrane (e.g., membrane fluidity). To test this, we quantified membrane fluidity in cells expressing either native TcpH or _EpsM_TcpH with and without α-linolenic acid using a fluorescent lipophilic pyrene-based probe (Figure S4D). Cells cultured with α-linolenic acid demonstrated elevated membrane fluidity, observed as a higher ratio of dimeric to monomeric pyrene probe (Figure S4D). We did not observe a change in membrane fluidity in WT cells cultured with linoleic acid (data not shown). These data demonstrate that the α-linolenic acid alters the biophysical properties of the membrane.

Poly-unsaturated fatty acids (PUFA), such as omega-3 fatty acids, influence lipid-ordered membrane domains within the cytoplasmic membrane of T-cells (73, 74). Lipid-ordered membrane domains, also called lipid rafts, are regions of the membrane enriched in saturated fatty acids, cholesterol (or, in some bacterial species, hopanoids), and proteins with specific TM domain qualities (typically long TM domain(s) and low surface area) (47, 55, 75). As a result, lipid ordered membrane domains tend to be thicker and less fluid than other areas of the membrane (47). n3-PUFA (i.e., omega-3 fatty acids) increase the size and stability of lipid-ordered membrane domains (47, 73, 74). We hypothesized that TcpP and TcpH molecules can associate within lipid-ordered membrane domains and that α-linolenic acid supplementation increases association of TcpP and TcpH molecules with the lipid-ordered membrane domain.

Lipid ordered membrane domains, also known as detergent resistant membranes (DRMs), were discovered due to their insolubility in Triton X-100 (52, 76). Triton X-100 has been used in both eukaryotic and prokaryotic organisms to isolate lipid ordered and disordered membrane domains (42–47). Thus, to test our hypotheses we used Triton X-100 to separate lipid ordered and lipid disordered membrane domains from cellular lysates.

Under Vir Ind conditions, TcpP and TcpH associate with Triton X-100 insoluble (TI; considered to be enriched with lipid ordered membrane domains) and Triton X-100 soluble membrane fractions (TS; considered to be enriched with lipid disordered membrane domains) (Figure 4AB). Supplementation with α-linolenic acid resulted in increases of both TcpP and TcpH in the TI fraction (Figure 4AB). Like TcpH, _EpsM_TcpH also associated with both the TI and TS membrane fractions (Figure 4CD). In contrast to native TcpH, there was no observable increase in _EpsM_TcpH levels in the TI fraction upon growth with α-linolenic acid (Figure 4CD). These data suggest that the native transmembrane domain of TcpH enables enhanced association with TI fractions (lipid ordered domains), and with the TcpP also in these fractions, after growth in α-linolenic acid.

**Figure 4:**
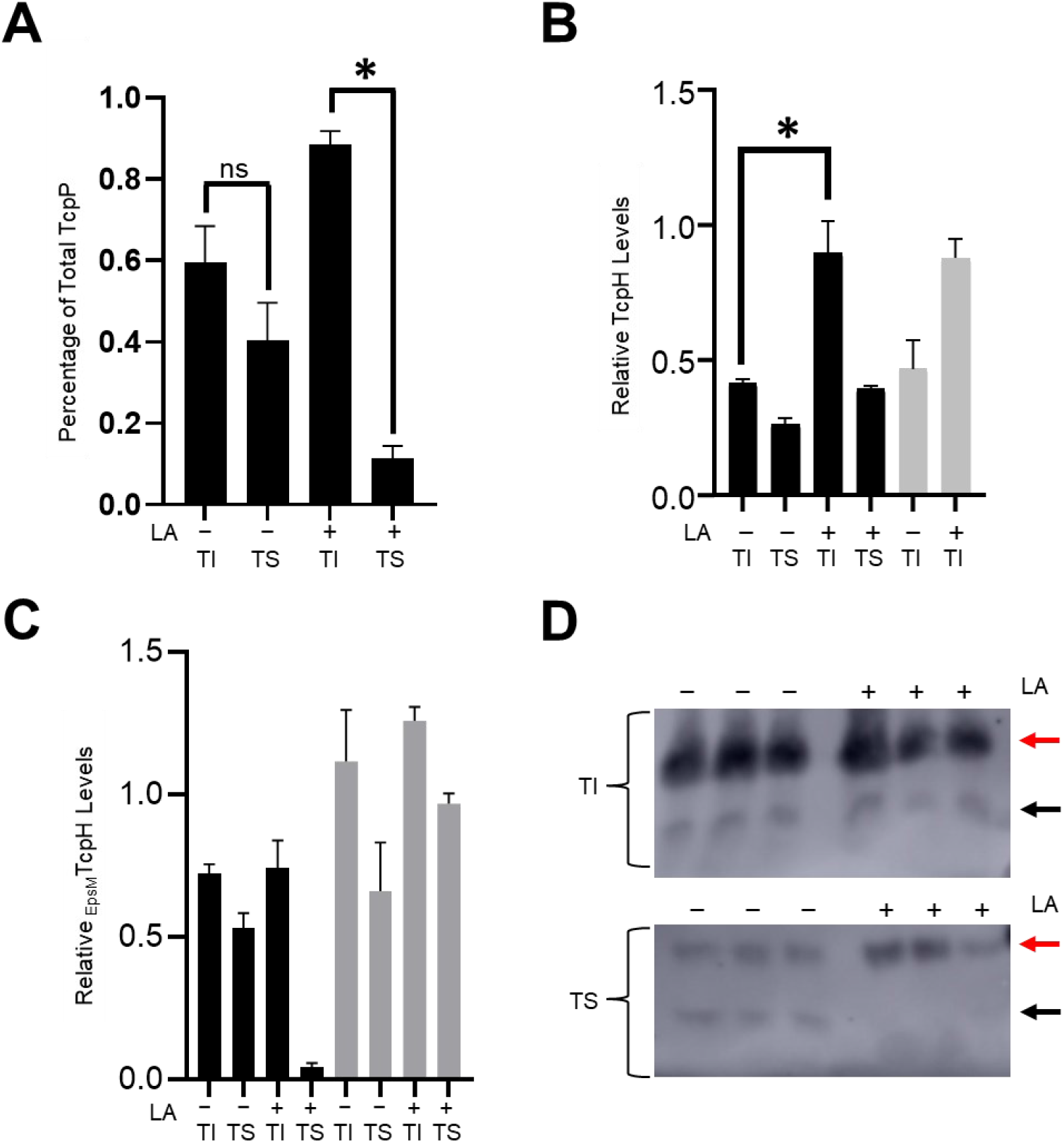
TcpP and TcpH abundance increases in detergent resistant membranes in the presence of α-linolenic acid. A) Percentage of total TcpP molecules within the Triton soluble (i.e., TS; lipid disordered) and Triton insoluble (i.e., TI; lipid ordered) fractions in WT cells. Percentage of TcpP within the TI and TS fractions was calculated by normalizing to the total amount TcpP in both the TI and TS fractions. B and C) Relative levels of TcpH and _EpsM_TcpH within the TI and TS membrane fractions. A-C) TcpP and TcpH levels were measured via densitometry using ImageJ. B and C) Black bars indicate TI and TS membrane fractions collected by spheroplast lysis, and gray bars indicate TI and TS samples collected using a gentle freeze thaw lysis. Cells that were cultured in α-linolenic acid (LA, 500µM) are indicated by +. TcpH and _EpsM_TcpH levels were normalized to a non-specific band (19KDa) that is equally distributed within TI and TS fractions (Figure S4E). D) Representative western blots of _EpsM_TcpH TI and TS membrane fractions. Black arrows mark the TcpH bands, and red arrows mark the non-specific band that serves as a loading control. Error bars represent the standard error of the mean. A two-tailed student’s T-test was used to determine statistical significance. * indicates a p-value less than 0.05, and ns indicates a lack of statistical significance.

Prior studies revealed that studying lipid ordered membrane domains with this biochemical method can yield dramatically different results with changes in detergent concentration and temperature (77). We thus performed similar experiments but used an alternative biochemical method to extract lipid ordered membrane domains. By altering the lysis method and the temperature at which cell lysis occurs, we observed the same trend of TI and TS association for TcpH and _EpsM_TcpH with and without α-linolenic acid present (Figure 4BC). We observed a shift in the percentage of TcpP molecules present in the TI and TS fraction (∼40% of TcpP molecules were present in the TI fraction and the remaining ∼60% was present in the TS) under Vir Ind conditions (Figure S4E). However, upon supplementation of α-linolenic acid to Vir Ind conditions, TcpP molecules maintained their higher localization to the TI fraction despite the change in our extraction method (Figure S4E). All told, these data suggest that enhanced RIP inhibition occurs due to increased association of both TcpP and TcpH with the TI fraction, and that the TM domain of TcpH drives this association with the TI fraction upon α-linolenic acid supplementation.

Excluding _EpsM_TcpH, it remained unclear if α-linolenic acid supplementation induced a general association of membrane proteins to the TI fraction. To test this, we quantified levels of a 19KDa non-specific membrane protein in TI and TS fractions with and without α-linolenic acid (Figure S4F). We observed no change in TI or TS abundance of this protein with α-linolenic acid supplementation (Figure S4F). These data indicate that α-linolenic acid supplementation does not induce a general association of proteins with the TI fraction. Furthermore, we also observed that with α-linolenic acid supplementation the TI fraction had a higher association of 16:0 fatty acids and lower association of 18:3 fatty acids than the TS fraction (Figure S4G). This is consistent with prior studies indicating that lipid ordered membrane domains are enriched with saturated fatty acids (53).

### TcpP and TcpH Interaction is critical for inhibition of RIP

Our data indicate that increased association of TcpP and TcpH molecules in the TI fraction results in enhanced RIP inhibition. The mechanism underlying this RIP inhibition remains unclear. Lipid-ordered membrane domains (which are also Triton insoluble) function as protein concentrators and thereby promote interaction between membrane localized proteins (49). We hypothesized that enhanced co-association within the TI fraction increased RIP inhibition due to direct interaction between TcpP and TcpH.

To test direct TcpP-TcpH interaction, we used a co-affinity precipitation approach. We genetically fused a His(6x)-Hsv or Hsv-His(6x) tag to the C-terminus and N-terminus, respectively, of TcpP, resulting in *tcpP-His-Hsv* and *Hsv-His-tcpP*. We could then extract TcpP from membrane fractions using NTA-Ni beads and identify TcpH and TcpP in elution fractions with α-TcpH and α-Hsv antibody. Proteins tagged at the amino-terminus are described with the tag noted first (e.g., Hsv-His-TcpP), while those tagged at the carboxy-terminus are described with the tag noted second (e.g., TcpP-His-Hsv).

First, we tested if both the N- and C-terminally-tagged proteins (Hsv-His-TcpP and TcpP-His-Hsv, respectively) function like native TcpP by measuring CtxB production after induction of the fusion proteins with arabinose under Vir Ind conditions. CtxB production was similar to that from cells expressing native TcpP, irrespective of which terminus the tag was placed (Figure S5).

Co-precipitation experiments indicated that the C-terminally-tagged TcpP could associate with TcpH, while the N-terminally-tagged TcpP could not (Figure 5AB). Physical interaction between the C-terminally tagged TcpP and TcpH also correlated to protection from RIP, as determined by assessing stability of the tagged proteins in cells expressing the first-site RIP protease Tsp but lacking the second protease YaeL. In such cells, the product of Tsp action on TcpP accumulates because the second-site protease YaeL is not present to eliminate it (26, 27). We observed greater accumulation of TcpP degradation intermediates (between 24KDa and 19KDa) in cells expressing N-terminally-tagged-TcpP compared to those expressing C-terminally-tagged TcpP (Figure 5C). The 24 kDa TcpP degradation intermediate from N-terminally-tagged TcpP is also observed in cells expressing native TcpP in the absence of TcpH (Figure 5CD). Considering that the N-terminally-tagged TcpP is sensitive to RIP even with TcpH present suggests a defect in its association with TcpH and its recognition by the RIP proteases. Despite this defect, N-terminally-tagged TcpP is capable of supporting WT CtxB production (Figure S5). We conclude this is the result of overexpression of N-terminally-tagged TcpP. Native expression of *tcpP* leads to accumulation of only TcpP* in a Δ*tcpH* Δ*yaeL* background (Figure 1E), but overexpression of *tcpP* in a Δ*tcpP* Δ*tcpH* Δ*yaeL* background yields both full length and TcpP* (Figure 5D). These data indicate that artificial elevation of TcpP levels, via overexpression, can outpace RIP.

**Figure 5:**
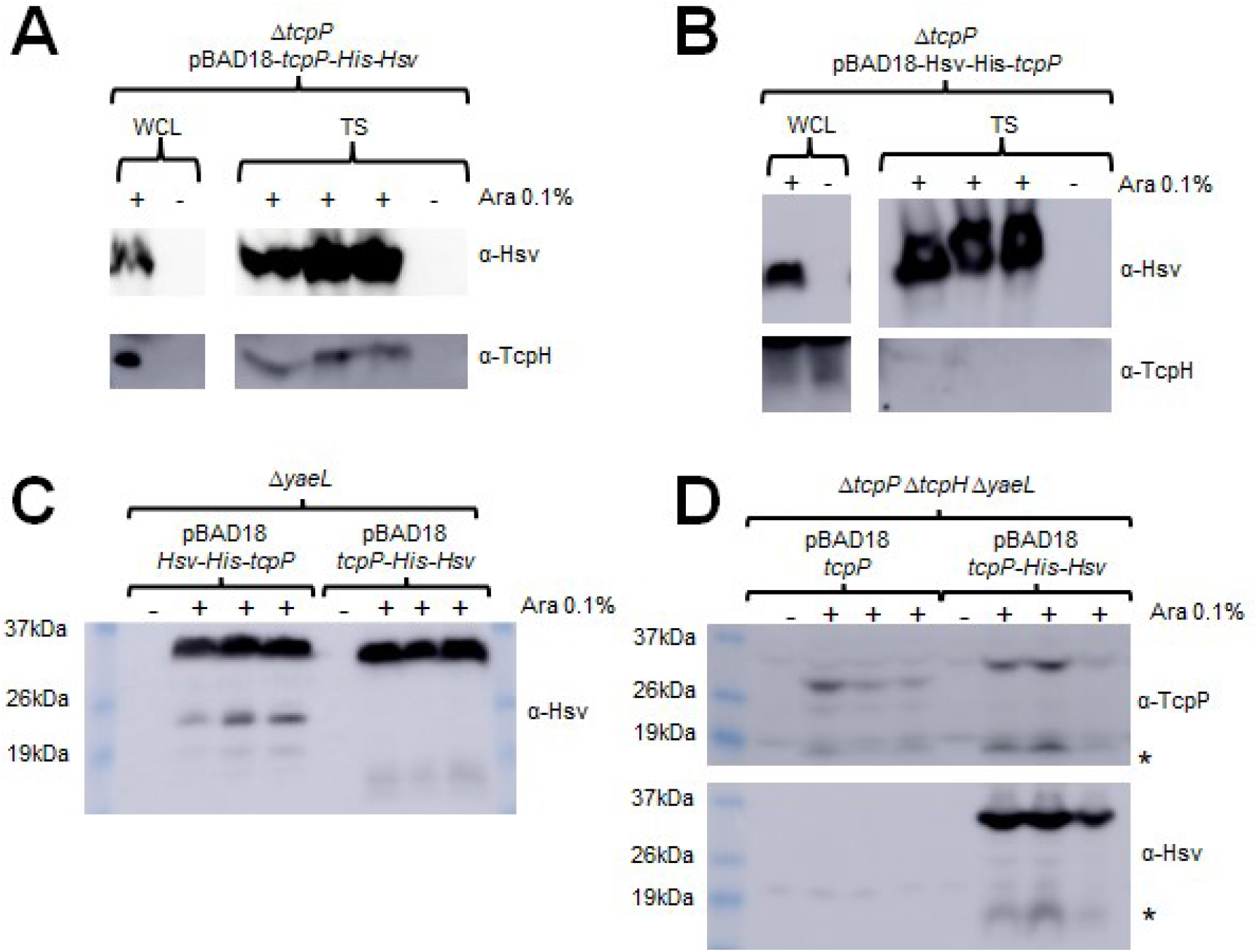
TcpP and TcpH interaction is critical for TcpH-dependent inhibition of RIP. A and B) Co-affinity precipitation of ectopically expressed *tcpP-His-HsV* (A), *Hsv-His-tcpP* (B). The data here represent three biological replicates. Triton soluble (TS). C) Ectopic expression of *Hsv-His-tcpP* and *tcpP-His-HsV* in Δ*yaeL* cells under virulence inducing conditions. Hsv-His-TcpP is more sensitive to RIP than TcpP-His-Hsv, as seen by accumulation of TcpP degradation intermediates between 26 and 19 kDa. A-C) *tcpP* constructs were all ectopically expressed from pBAD18 using arabinose (Ara 0.1% w/v). + indicates arabinose was added to the culture. Samples presented here represent three biological replicates.

Our data indicate that TcpP-His-Hsv is less sensitive to RIP in the presence of TcpH. Prior studies have demonstrated that modification of the C-terminus of TcpP can lead to TcpH-independent resistance to RIP (78). To determine if the addition of His-Hsv to the C-terminus of TcpP promotes resistance to RIP independent of TcpH we expressed *tcpP-His-Hsv* and *tcpP* in a Δ*tcpP* Δ*tcpH* Δ*yaeL* background. We observe TcpP* accumulation in both *tcpP* or *tcpP-His-Hsv* expressing cells (∼17KDa) (Figure 5D). These data show that addition of His(6x)-Hsv to the C-terminus of TcpP does abrogate the need for TcpH to protect TcpP-His-Hsv from RIP (Figure 5D). In summary, our data indicates that TcpP and TcpH interact and that this interaction is important for inhibition of RIP of TcpP.

It remains unclear why Hsv-His-TcpP is unable to interact with TcpH. Single-molecule tracking studies indicate that TcpP may be sensitive to RIP while interacting with the *toxT* promoter (78). The Hsv tag is enriched with negatively charged amino acids (Hsv amino acid sequence: QP**E**LAP**ED**P**ED**). Given that DNA has an intrinsic negative charge, addition of Hsv-His(6x) to the N-terminus of TcpP may promote a conformation similar to the conformation that TcpP molecules adopt when actively interacting with DNA. This hypothesis requires additional experiments to test.

### Miltefosine, a lipid raft busting drug, functions synergistically with α-linolenic acid

*Staphylococcus aureus* relies on lipid ordered membrane domains to recruit and promote oligomerization of flotillin, which in turn promotes antibiotic resistance (45). Miltefosine, a drug used to treat Leishmaniasis and certain types of cancers, inhibited flotillin association with lipid ordered membrane domains in *S. aureus* (45, 79). Our data indicate that α-linolenic acid enhances *toxT* expression by promoting association of TcpP and TcpH molecules within lipid ordered membrane domains. We hypothesized that miltefosine treatment would inhibit TcpH dependent enhanced RIP inhibition in the presence of α-linolenic acid. Instead, we observed that miltefosine alone functioned like α-linolenic acid (Figure S6A). Treatment with both miltefosine and α-linolenic acid resulted in a ∼7-fold increase in TcpP levels relative to Vir Ind conditions (Figure S6B). Our data also demonstrate that miltefosine also promoted association of TcpP molecules with the TI fraction like α-linolenic acid (Figure S6C). Miltefosine did not promote *toxT* transcription in Δ*tcpH* and _EpsM_TcpH cells (Figure S6A).

Taken together, these data indicate that miltefosine functions synergistically with α-linolenic acid to increase levels of TcpP in *V. cholerae* and is not effective at inhibiting lipid ordered domain formation in *V. cholerae*. Miltefosine is known to associate with lipid ordered domains and requires lipid ordered domains to enter cells (80, 81). Also, miltefosine increases membrane fluidity (82). Other n3-polyunsaturated fatty acids, like α-linolenic acid, are also capable of increasing membrane fluidity, and they have been shown to drive aggregation and stabilization of lipid ordered membrane domains (47, 73, 74). Given that miltefosine and α-linolenic acid function synergistically to promote TcpH-dependent antagonism of RIP, these data suggest that α-linolenic acid promotes lipid ordered domain aggregation, and thereby increases lipid ordered domain size in *V. cholerae* cells.

## Discussion

Canonical RIP systems act by releasing an anti-sigma factor from the cytoplasmic membrane to influence gene expression (28, 83). Membrane localized transcription regulators (MLTRs), in addition to TcpP and ToxR, are sensitive to RIP (e.g., CadC) (84). However, RIP of MLTRs, such as TcpP, results in their inactivation, typically leading to decreased gene expression. The fundamental mechanisms of RIP for TcpP are understood, in terms of the primary proteases that work in the two-step pathway (26, 27), but the regulatory mechanisms influencing these activities have been less well studied. It is clear that TcpH is essential to inhibit RIP of TcpP, and that its ability to protect TcpP from RIP changes in response to temperature and pH (25–27). ToxR is a well-studied MLTR, similar to TcpP and is sensitive to RIP (85, 86). ToxR is protected from RIP by ToxS, a single pass transmembrane protein analogous to TcpH, via direct interaction (12, 87). Prior work indicates that: i) ToxR undergoes RIP during late stationary phase (i.e., alkaline pH and nutrient limiting conditions); ii) ToxS antagonizes RIP of ToxR via direct interaction; and iii) deoxycholate increases interaction between ToxR and ToxS (88-91). As is understood about ToxR, our data indicate that RIP of TcpP is inhibited by direct interaction with TcpH. Our data indicate that α-linolenic acid, a host dietary fatty acid, plays a role in inhibiting RIP by increasing the local concentration of TcpP and TcpH within detergent resistant membranes (DRM) (i.e., lipid ordered membrane domains). Whether this fatty acid plays any role in ToxR RIP inhibition remains to be examined.

α-Linolenic acid is an essential omega-3 fatty acid used to synthesize arachidonic and docosahexaenoic acid humans and mice (92, 93). α-Linolenic acid is acquired via dietary supplementation and is present in milk, meats, dairy products, soybean oil, and plant seeds (94-100). It is considered a beneficial dietary fatty acid as it is a precursor to omega-3, omega-6, and conjugated α-linolenic acids, and has health benefits ranging from anti-carcinogenic, anti-atherogenic, anti-inflammatory, improved memory, and anti-diabetic activity (101-109). *V. cholerae* uses exogenous long-chain fatty acids, such as α-linolenic acid, to remodel its phospholipid composition (61, 62). Long-chain fatty acids are transported across the outer membrane by FadL into the periplasmic space where FadD covalently modifies the fatty acids by adding an acyl-CoA group, resulting in formation of long-chain fatty acyl-CoA (LCFA-CoA) (57–60). LCFA-CoAs then bind to FadR, the principal regulator of fatty acid biosynthesis in *V. cholerae*, resulting in a conformational change inhibiting FadR from binding to DNA (110-112). This leads to decreased biosynthesis of unsaturated fatty acids (i.e., decrease in *fabAB* expression) and increased expression, due to a lack of repression by FadR, of genes required for transport, activation, and beta-oxidation of long-chain fatty acids (i.e, *fadL, fadD, fadBA, fadE*, and *fadH*) (110-112). FadL into the periplasmic space where FadD covalently modifies the fatty acids by adding an acyl-CoA group, resulting in formation of long-chain fatty acyl-CoA (LCFA-CoA) (57–60). LCFA-CoAs then bind to FadR, the principal regulator of fatty acid biosynthesis in *V. cholerae*, resulting in a conformational change inhibiting FadR from binding to DNA (110-112). This leads to decreased biosynthesis of unsaturated fatty acids (i.e., decrease in *fabAB* expression) and increased expression, due to a lack of repression by FadR, of genes required for transport, activation, and beta-oxidation of long-chain fatty acids (i.e, *fadL, fadD, fadBA, fadE*, and *fadH*) (110-112). FadL into the periplasmic space where FadD covalently modifies the fatty acids by adding an acyl-CoA group, resulting in formation of long-chain fatty acyl-CoA (LCFA-CoA) (57–60). LCFA-CoAs then bind to FadR, the principal regulator of fatty acid biosynthesis in *V. cholerae*, resulting in a conformational change inhibiting FadR from binding to DNA (110-112). This leads to decreased biosynthesis of unsaturated fatty acids (i.e., decrease in *fabAB* expression) and increased expression, due to a lack of repression by FadR, of genes required for transport, activation, and beta-oxidation of long-chain fatty acids (i.e, *fadL, fadD, fadBA, fadE*, and *fadH*) (110-112).

Utilization of exogenous fatty acids remodels the phospholipid bilayer in *Vibrio spp*. (61, 62, 113) and has an impact on pathogenicity, motility, and antibiotic resistance via unknown mechanisms (62). Our work demonstrates that: i) *toxT* expression is enhanced in the presence of α-linolenic acid; ii) TcpP levels are significantly elevated in the presence of α-linolenic acid; iii) the *tcpP* transcript level is not increased with exogenous α-linolenic acid; iiiv) TcpP and TcpH avidly associate within detergent resistant membranes (DRM; hypothesized to be lipid-ordered domains) in the presence of α-linolenic acid; v) TcpP-TcpH interaction is important for inhibition of RIP; And vi) enhanced *toxT* expression in the presence of α-linolenic acid is dependent on co-association of TcpP and TcpH in the DRM membrane fraction.

Our data support a model where, once present in the gastrointestinal tract (GI), *V. cholerae* cells take up and incorporate α-linolenic acid, present in the GI tract of infant mice (114), into phospholipids, thereby altering the composition of the cytoplasmic membrane. This influences TcpH and TcpP molecules to increase their association with lipid ordered membrane domains via an unknown mechanism. n-3 polyunsaturated lipids (i.e., omega-3 fatty acids) are known to increase lipid ordered domain size in eukaryotes by promoting aggregation of existing lipid ordered membrane microdomains (73, 74). As lipid ordered membrane domains are known to be relatively small in size (6-200 nm) (49, 51), we hypothesize that this leads to an increase in the local concentration of TcpP and TcpH molecules thereby allowing TcpH to enhance RIP inhibition of TcpP via increased interactions with TcpP (Figure 6).

**Figure 6:**
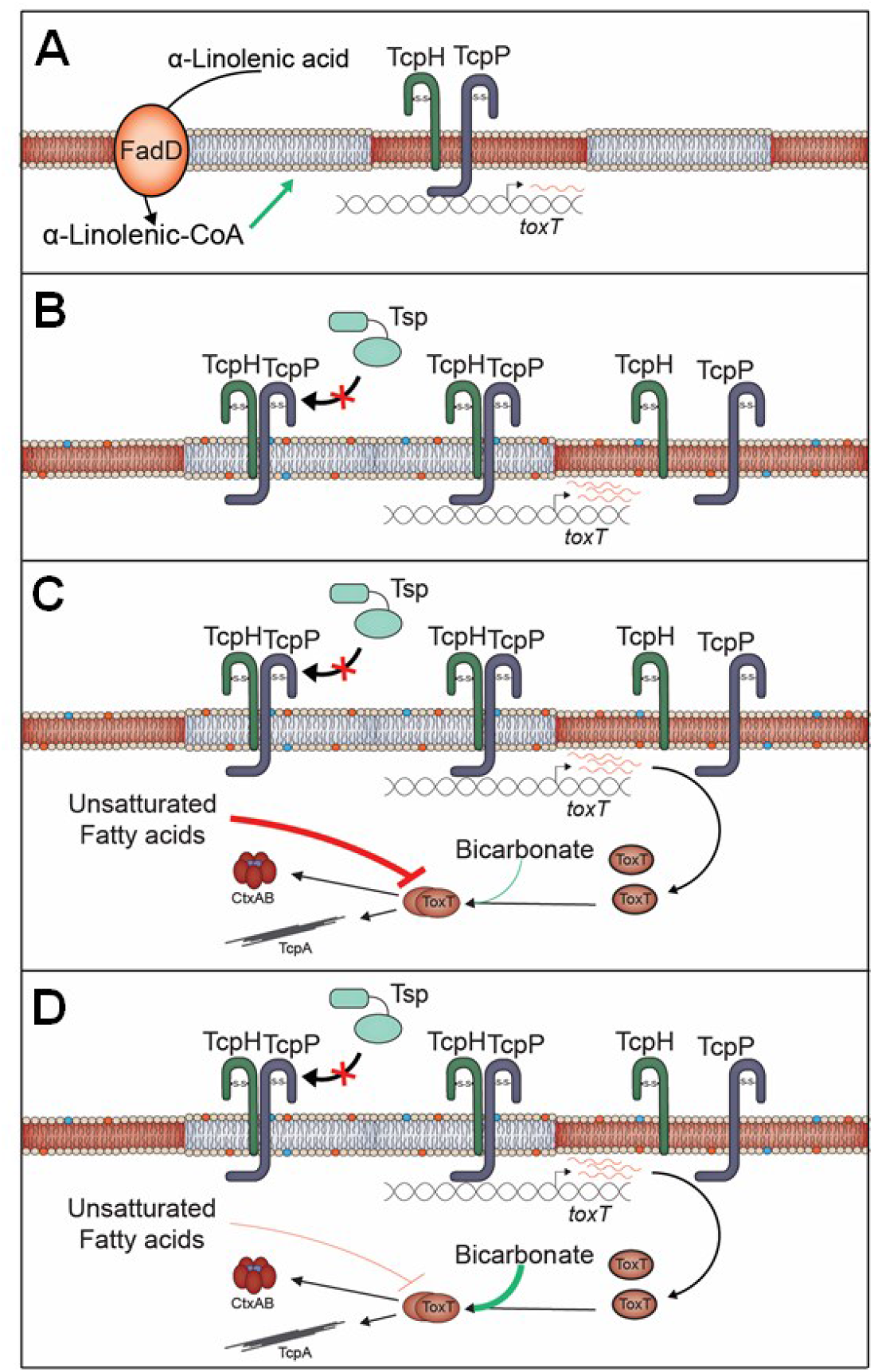
α-Linolenic acid stimulates co-association of TcpP and TcpH within detergent resistant membranes promoting TcpH-dependent inhibition of RIP. A) Under virulence inducing (Vir Ind) conditions TcpP and TcpH molecules are associated within Triton insoluble (blue; TI) and Triton soluble (red; TS) membrane domains. B) In the presence of exogenous α-linolenic acid, *V. cholerae* cells uptake α-linolenic acid (via VolA, FadL) and utilize it directly for phospholipid remodeling via addition of coenzyme A (CoA), via FadD (57–60). This leads to changes in the overall phospholipid profile of *V. cholerae*, indicated by the blue and orange phospholipids (58–61). Under these conditions, a majority of TcpP and TcpH molecules transition to the TI membranes leading to enhanced inhibition of RIP by TcpH. The net result of α-linolenic acid supplementation is an increase in *toxT* transcription, indicated by an increase in red *toxT* mRNA. C) In the lumen of the gastrointestinal tract, ToxT activity is thought to be inhibited by unsaturated fatty acids, and thus inhibits ToxT-dependent virulence factor expression (63, 64, 116, 122–124). D) Near the surface of epithelial cells, bicarbonate is actively secreted from intestinal epithelial cells and, as such, the concentration of bicarbonate is elevated, including the crypt of intestinal villi (122–124). As Bicarbonate is known to stimulate ToxT activity, ToxT-dependent virulence gene expression is thought to be stimulated near epithelial cells (122-124).

Previous studies have investigated the role of exogenous fatty acids on the pathogenesis of *V. cholerae*. These concluded that FadD is required for wild-type *toxT* expression through a mechanism involving its effect on TcpP levels (115, 116). These prior publications support our model as accumulation of α-linolenic acid in the periplasmic space or within the cytoplasmic membrane, due to loss of *fadD*, results in a reduction in TcpP levels, rather than an increase (115, 116). This work indicates that free α-linolenic acid (I.e., not incorporated in phospholipids) within the periplasmic space, cytoplasm, or within the cytoplasmic membrane, does not promote TcpH mediated inhibition of RIP. In conjunction with the data presented here, this indicates that α-linolenic acid needs to be incorporated into the cytoplasmic membrane as a phospholipid to have an effect on TcpH function.

Lipid ordered and lipid disordered membrane domains were discovered due to the insolubility of the lipid ordered membrane domain (initially referred to as detergent resistant membranes) in Triton X-100 and other non-ionic detergents (52, 76). This biochemical property has been used to separate lipid ordered (DRM) and lipid disordered (DSM) membrane domains in many Eukarya and Bacteria, including Gram-negative and Gram-positive bacteria (42–47). Data generated from the biochemical-based separation of lipid ordered and lipid disordered membrane domains has been verified by alternative methods (e.g., fluorescent microscopy, single-molecule tracking, and synthetic membrane vesicles) (117). Due to a lack of literature on lipid ordered and lipid disordered membrane domains in *V. cholerae*, we performed additional experiments to determine if our biochemical extraction method faithfully enriched for lipid ordered membrane domains and lipid disordered membrane domains within the DRM and DSM (I.e., TI and TS) respectively. In the presence of α-linolenic acid, we found that the TI fraction had a higher association of 16:0 fatty acids and a lower association of 18:3 fatty acids compared to the TS fraction (Figure S4F). These characteristics are consistent with lipid ordered membrane domain and suggest that the TI and TS fractions presented here are enriched in lipid ordered and lipid disordered membrane domains respectively.

Transmembrane domain length and surface area are major factors in determining the preference of a protein for lipid ordered (enriched with proteins having longer TM domain and low surface area) or lipid disordered (enriched with proteins having shorter TM domain and high surface area) membrane domains (118). We demonstrated that TcpH and TcpP increase localization within DRM domains in the presence of α-linolenic acid while _EpsM_TcpH does not (Figure 4). _EpsM_TcpH has a shorter TM domain than TcpH (20 amino acids vs 22 amino acids) and a higher overall surface area (108 Å^2^ vs 92 Å^2^). Thus, we hypothesize that the TM domain properties of _EpsM_TcpH molecules inhibit its transition from the TS fraction to the TI fraction in the presence of α-linolenic acid. Alternatively, it is also possible that TcpH, and not _EpsM_TcpH, undergoes post-translational modification (e.g., palmitoylation) within its TM domain. We view this as unlikely as TcpH is not predicted to have a palmitoylation site within its TM domain. In addition, it also appears that the surface area of the transmembrane domain of TcpP influences its function. Prior analysis of TcpP transmembrane domain revealed that mutation of L152 and W162/S163 with alanine (which reduces the overall surface area of the transmembrane domain) increased *toxT* expression (119). It remains unclear why these mutations increase TcpP function, but given the data presented here, it is possible that TcpPL152A and TcpP W162A/S163A may have a greater propensity than TcpP, in the absence of α-linolenic acid, to associate within DRMs (i.e., lipid ordered membrane domain).

Based on our data here and other literature, we hypothesize that phospholipid remodeling of *V. cholerae* occurs in the lumen during the initial stages of infection. Our data suggests that this remodeling promotes TcpH mediated inhibition of RIP and promotes *toxT* transcription. However, unsaturated fatty acids can also inhibit degradation and activity of ToxT (63, 64, 120). This likely prevents premature expression of TCP which is known to stimulate microcolony formation and thereby could inhibit penetration of the mucus layer (121). Bicarbonate present at high concentrations at the surface of epithelial cells, competes with unsaturated fatty acids to activate ToxT once *V. cholerae* reaches the surface of epithelial cells, its primary site of infection (122– 124). There is also evidence that bicarbonate represses *toxT* transcription (123). This indicates that expression of *toxT*, stimulated by enhanced RIP antagonism, during early infection (i.e., the lumen) is critical for *V. cholerae* to cause disease. This adds a new level of regulation to the ToxR regulon and another dietary host factor that modulates *toxT* expression in *V. cholerae*. α-Linolenic acid represents the first *in vivo* signal that modulates RIP of TcpP, and, to the best of our knowledge, the first evidence that lipid ordered and lipid disordered membrane domains exist in *V. cholerae*. The data presented here further expands our knowledge of the complex virulence regulatory cascade in *V. cholerae*.

## Materials and Methods

### Bacterial culture conditions

All *V. cholerae* strains used in this study were of the classical biotype (0395) (See Table S1 for a complete list of bacterial strains). Unless otherwise stated *Escherichia coli* and *V. cholerae* were grown at 37°C in Luria-Bertani (LB; 10 g tryptone, 5 g yeast extract, and 5 g NaCl per liter) with vigorous shaking (210 rpm). LB was prepared as previously described (125). To stimulate virulence factor production, *V. cholerae* strains were subcultured, to an O.D. of 0.01, from an overnight LB culture and grown under virulence inducing conditions (Vir Ind; 30°C, LB pH 6.5 ∓0.5, and 110 rpm) or non-virulence inducing conditions (non-Vir Ind; 37°C, LB pH 8.5 ∓0.5, and 210 rpm). Media used for both Vir Ind and non-Vir Ind were sterilized using 1L 0.22 µm vacuum filtration units (Sigma) after pH adjustment.

*ex-vivo* mouse fecal experiments with sterile and non-sterile mouse fecal media were conducted aerobically at 37°C in 48 well plates (Sigma) with shaking (210 rpms). Sterile mice fecal samples were collected from C57 Black female mice on 4 separate days and stored at -80°C. After collection mice fecal samples were homogenized, via mortar and pestle, and then suspended in M9 minimal media. The final concentration of mice fecal media was 9% w/v. The mice fecal media was then spun down (2450xg for 10 min) to remove insoluble material. The supernatant was collected, and filter sterilized using a 0.45 µM syringe filter (Sigma). Non-sterile mice fecal samples were collected from C57 Black female mice on three separate days. Mice fecal matter was directly resuspended in M9 media to a final concentration of 9% w/v. Mice fecal media was then incubated at room temperature for 1 hour while shaking on a table top shaker. Mice fecal media was spun down (2450xg for 10 min). The supernatant was collected and used directly for the growth curve. *V. cholerae* cell density was determined by counting CFU’s on LB agar plates supplemented with streptomycin. Microbiota in mice fecal matter were not found to be resistant to streptomycin.

To test if crude bile (Ox gal, Sigma Aldrich), as well as components of crude bile, we opted to pretreat all *V. cholerae* strains under Vir Ind conditions before exposing cells to these additional factors. *V. cholerae* cells were subcultured from overnight cultures to an optical density of 0.01 in 100 ml of LB pH 6.5 in a 250 ml Erlenmeyer flask. *V. cholerae* strains were grown for 4 hours under Vir Ind conditions, centrifuged (2450 X g 15 minutes), resuspended in 0.8 ml LB. 200 µl of resuspended cells were transferred to 50 ml of fresh Vir Ind media in 125 ml erlenmeyer flasks. The remaining 200 µl of cells were lysed and analyzed via western blot. A maximum of 4 different conditions were tested per strain per biological replicate due to limited incubator space. The following were supplemented to Vir Ind media: crude bile (CB; final concentration 0.4% v/v), α-linolenic acid (LA; final concentration 500µM-500nM), palmitic acid (PA; final concentration 500µM), stearic acid (SA; final concentration 500µM), purified bile salts cholate and deoxycholate (PB; final concentration 100µM). All compounds were purchased from Sigma Aldrich. CB and PB were solubilized in Vir Ind media and filter sterilized (0.22 µM; Sigma) before addition to Vir Ind. LA and PA were dissolved in Dimethyl sulfoxide (DMSO) and then added to Vir Ind media. LA and PA sterility were confirmed by spreading 100µl of DMSO solubilized LA and PA on LB agar plates (data not shown).

Unless otherwise stated, antibiotics were used at the following concentrations: ampicillin (100 µg/ml), chloramphenicol (30 µg/ml), streptomycin (100 µg/ml), and cerulenin (10 µg/ml). Overexpression of constructs by pBAD18 was induced by culturing strains in LB containing 0.1% arabinose.

### Plasmid construction

Briefly, DNA fragments 500 bp upstream and downstream of the target gene were amplified using Phusion high-fidelity polymerase (Thermo Scientific) (see Table S2 for list of primers used). Insert fragments containing desired mutations were connected by splicing via overlap extension PCR. Plasmid vectors (pKAS32 and pBAD18) were isolated from bacterial strains using the Qiagen Miniprep kit. Plasmid vectors were then digested with KpnI-HiFi and XbaI (New England BioLabs) at 37°C for 2 hours. Insert and vector fragments were then added to Gibson assembly master mix (New England BioLabs) and incubated at 50°C for 30 minutes. Plasmids were then introduced to *E. coli* ET12567 *ΔdapA* (λpir +) by electroporation. pKAS32 plasmids were then transferred to *V. cholerae* strains via mating on LB agar plates at 30°C overnight. pBAD18 plasmids were introduced into *V. cholerae* strains via electroporation.

### Mutant construction

Mutants were constructed as previously described (126). *V. cholerae* harboring pKAS32 derivatives were grown in 2 ml LB for 2 hours at 37°C. Streptomycin was then added to cultures to a final concentration of 2500 µg/ml and incubated for an additional 2 hours. After a total of 4 hours of incubation, 20 µl of culture was spread on LB agar plates containing streptomycin (2500 µg/ml) and incubated at 37°C overnight. Colonies that were resistant to streptomycin were screened via colony PCR to confirm presence of the desired mutation. Genomic DNA was then isolated from potential mutants and the region of interest was then amplified via PCR and validated by sequencing (GeneWiz).).

### Growth curves

*V. cholerae* strains were subcultured from an overnight culture to a final optical density (600 nm) of 0.01 in 200 µl of virulence inducing media, LB, or M9 minimal media (supplemented with 0.05% glucose) per well of a 96 well plate. The plate was then incubated at 30°C or 37°C in a SPECTROstar Omega plate reader (BMG LABTECH), with shaking and optical density measurements every 30 minutes.

### Western blots

After whole cell lysis, the total protein concentration of each sample was measured via Bradford assay (Sigma Aldrich). Samples were subsequently diluted to a final concentration of 0.5 µg total protein/µl. All SDS page gels contained 12.5% acrylamide and were run at 90-120 volts for 1.5 hours. Proteins were transferred to nitrocellulose membranes using a semi dry electroblotter (Fisher Scientific) overnight at 35 mA or for 2 hours at 200mA. Membranes were blocked with 15 ml of blocking buffer (5% non-fat milk, 2% bovine serum albumin, 0.5% Tween-20, in Tris-buffered saline) for 1 hour at room temperature. Primary antibodies were diluted in 5% non-fat milk and Tris-buffered saline (α-TcpH 1:500, α-TcpP 1:1,000, and α-TcpA 1:100,000) and incubated with the membranes for 1 hour at room temperature. Membranes were washed three times for 5-15 minutes with Tris-buffered saline. Secondary antibodies (Sigma Aldrich) were diluted in 5% non-fat milk in Tris-buffered saline (Goat anti-Rabbit IgG-HRP 1:2,000) and incubated as before. Membranes were washed three times for 5-15 minutes with Tris-buffered saline and then incubated with SuperSignal HRP Chemiluminescence substrate (Thermo Fisher). Membranes were imaged with an Amersham Imager 600.

### Enzyme Linked-Immunosorbent Assay (ELISA)

ELISAs were performed as previously described (127). *V. cholerae* cells were subcultured from overnight cultures to an optical density of 0.01 in 10 ml of LB pH 6.5. Cultures were incubated at 30°C for a total of 24 hours. Cells were collected by centrifugation at 2450X g for 15 minutes. 1 ml of culture supernatant was collected and the remaining supernatant was discarded. All steps of EILSA were performed at room temperature. 10 µl of culture supernatant was added to 140 µl PBS-T (phosphate buffered saline, 0.05% Tween-20, 0.1% BSA) in row A of plates coated with GM1 (monosialotetrahexosylganglioside). Samples were diluted (1:3) down each column and incubated at room temperature for 1 hour. Plates were then washed with PBS-T three times. Primary (α-CtxB 1:8000, Sigma Aldrich) and secondary antibody (Goat anti-Rabbit IgG-HRP 1:5,000, Sigma Aldrich) were diluted in PBS-T. 100 µl of diluted antibody was added to each well and incubated for 1 hour at room temperature. Plates were again washed with PBS-T as before. 100 ul of TMB (3,3’,5,5’-tetramentylbenzidine, Sigma) was added to each well and incubated for 5-10 minutes. The reaction stopped by addition of 100 µl of 2M sulfuric acid and the optical density (450 nm) was measured for each well using SPECTROstar Omega plate reader (BMG LABTECH).

### Infant Mouse Colonization

Infant mouse colonization experiments were performed as previously described (128, 129). The Institutional Animal Care and Use Committee at Michigan State University approved all animal experiments (PROTO201900421). 3-6 day old male and female Infant mice (CD1, catalog #: 022CD1) were purchased from Charles River (Wilmington, MA). Infant mice were transported with a female adult mouse. Upon arrival, infant mice were separated from the adult female mouse, 2 hours prior to infection, and all adult female mice were euthanized. Infant mice were randomly assigned into infection groups. 70% ethanol was used to sterilize gloves, tubing, and surfaces between groups. The number of mice used per group is indicated in the figure or figure legend. Briefly, three- to six-day old CD-1 mice were orogastrically inoculated with ∼1×10^6^ or ∼1×10^8^ bacterial cells after 2 hours of separation from their mothers. Infant mice were kept at 30°C in sterile bedding and euthanized either 18 hours or 21 hours after infection. Mouse intestines (small and large) were weighed in 3 ml PBS and homogenized. For fluid accumulation studies, infant mice were weighed prior to collection of mouse intestine, and mouse intestines were weighed after blotting on absorbent paper. Homogenates were then serially diluted in PBS, spread on LB plates containing streptomycin, and incubated at 37°C overnight.

### Real-time quantitative PCR (RT-qPCR)

RT-qPCR experiments were performed as previously described (130). RNA was preserved by resuspending *V. cholerae* cells in 1 ml of Trizol (Sigma Aldrich) and then extracted from cells using an RNEasy kit (Qiagen) according to manufacturer’s instructions. RNA was then treated with Turbo DNase for 30 minutes at 37°C. After DNase treatment, RNA quality was determined by detection of large and small ribosomal subunits via 2% agarose gel. RNA quantity was then measured using a Nanodrop spectrophotometer (Thermo Scientific). cDNA was generated from DNase treated RNA using Superscript III reverse transcriptase (Thermo Scientific) as previously described (130). 5 ng of cDNA was used with SYBR green master mix (Applied Biosystems) to perform the RT-qPCR. *recA* was used as a housekeeping gene of reference to calculate the threshold values (ΔΔC_T_) (131). See Table S2 for primers.

### β-Galactosidase activity assay

*V. cholerae* cells were subcultured from overnight cultures to an optical density of 0.01 in 50 ml of LB pH 6.5. *V. cholerae* strains were grown for 4 hours under Vir Ind conditions. Following incubation cultures were centrifuged (2450 X g 15 minutes), resuspended in 1 ml LB, and then 200 µl of the culture resuspension was transferred to fresh media (Vir Ind, Vir Ind supplemented with crude bile/ cholate and deoxycholate (purified bile)/ α-linolenic acid, or non-Vir Ind). Cultures were grown for an additional 4 hours under their indicated condition. At the indicated time point (4 hours or 8 hours) 1.5 ml of culture was removed, centrifuged (4000 X g 15 minutes), and resuspended in 1 ml of Z-buffer (Na_2_HPO_4_ 60mM, NaH_2_PO_4_ 40mM, KCl 10mM, MgSO_4_ 1mM, β-mercaptoethanol 50mM, pH7.0). β-galactosidase activity and Miller units were determined as previously described (132).

### Subcellular Fractionation

Cells were fractionated following the Tris-sucrose-EDTA method (200mM Tris-HCl pH 8.5, 500mM sucrose, 1mM EDTA, pH 8.0) (133). *V. cholerae* cells were subcultured from overnight cultures to an optical density of 0.01 in 50 ml of LB pH 6.5. After 2 hours of incubation, plasmids were induced by the addition of arabinose (final concentration of 0.1%) at 30°C with mild shaking (110 rpm), and then cultured for an additional 5 hours. All steps of the fractionation procedure were performed on ice as follows (133). Spheroplast fractions (i.e., cytoplasm and the cytoplasmic membrane) were resuspended in 500 µl 0.45% NaCl. To lyse the spheroplasts 50 µl of 10% SDS were added and samples were then boiled for 5-10 minutes. Periplasmic fractions were concentrated using trichloroacetic acid (TCA) (133, 134). Pelleted whole cells were resuspended in 50-200 µl of resuspension buffer (50mM Tris-HCl, 50mM EDTA, pH 8.0). Cells were then lysed by the addition of lysis buffer (10mM Tris-HCl, 1% SDS) and boiled for 5-10 minutes. All fractions were stored at - 20 °C until use.

Soluble and insoluble fractionation of *V. cholerae* cells was performed as described by Miller *et. al*., with modifications (11). Initial steps of the Tris-sucrose-EDTA extraction were followed regarding growth and collection of *V. cholerae* cells. Following collection, cells were resuspended in 10 ml of lysis buffer (10 mM Tris HCl pH 8.0, 750mM sucrose, EDTA-free protease inhibitor, 2mM EDTA, 50 µg/ml lysozyme, 10 U/ml DNase 1) and incubated on ice for 20 minutes. Cells underwent two rounds of lysis via French press (7,000-10,000 psi). Cellular debris was removed by centrifugation (1200 X g for 10 minutes) and supernatant was retained. Insoluble (i.e., the inner and outer membrane) and soluble fractions were separated by ultracentrifugation (100,000 x g for 2 hours at 4°C). The pellet, containing the membrane fraction, was collected and resuspended in 500 µl 5mM EDTA and 25% sucrose. The insoluble membrane fraction underwent a second round of ultracentrifugation and was then collected. All samples were stored at -80°C until further use.

### Fatty Acyl Methylester (FAME) analysis

Analysis of fatty acids from whole *V. cholerae* cells was done as previously described (135). Briefly, *V. cholerae* cells were grown with and without linolenic acid (500 µM) as described in the section below. Cells were collected by centrifugation (2450 X g 15 minutes) and then washed with PBS. Cells were then lysed via addition of 300 μl of extraction solvent (composed of methanol, chloroform and formic acid [20:10:1, v/v/v]). After lipids were extracted the Fatty Acyl Methylester (FAME) reactions were carried out as described (135). After the FAME reactions, fatty acid content was measured via Gas-Liquid Chromatography using a DB-23 column (agilent, part number: 122-2332). Molar values of each peak were then normalized to an internal standard (15:0) to calculate the total molar percentage of each fatty acid detected.

### Membrane Fluidity

Membrane fluidity was measured as previously described (136) using a membrane fluidity kit that quantifies the fluorescence of a lipophilic dye (Pyrenedecanoic acid) (Abcam). Pyrene decanoic acid exists in monomeric and dimeric states within membranes. Dimerization of pyrene decanoic acid occurs in areas of low fluidity (or low viscosity) and results in a change in its emission spectra. Thus, the ratio of dimeric (excimer state) and monomeric pyrene decanoic acid can be used to quantify membrane fluidity. Briefly, WT and _EpsM_TcpH cells were subcultured from overnight cultures to an optical density of 0.01 in 100 ml of LB pH 6.5 in a 250 ml Erlenmeyer flask. *V. cholerae* strains were grown for 4 hours under Vir Ind conditions, centrifuged (2450 X g 15 minutes), resuspended in 0.8 ml LB. 200 µl of resuspended cells were transferred to 50 ml of fresh Vir Ind media supplemented with ethanol (3%w/v), benzyl alcohol (20mM), DMSO (1% w/v), or α-linolenic acid (500 µM) in 125 ml erlenmeyer flasks. Cultures were incubated for an additional 4 hours under Vir Ind or Non-Vir Ind conditions. After incubation cells were collected from 1 ml of culture via centrifugation (2450 X g 15 minutes) and resuspended in 500 µl LB. Cells were incubated with the fluorescent lipid reagent (10 µM final concentration) for 30 minutes at room temperature (∼23°C) while shaking. Cells were then washed twice with LB and fluorescence (excitation, 350 nm, and emission, 400 nm and 470 nm) was quantified for each sample. After subtracting the background fluorescence, the fluorescence ratio was calculated for each sample by dividing the excimer (470 nm) by the monomer (400 nm) fluorescence. Unlabeled cells and non-Vir ind conditions were used as negative controls. Ethanol and benzyl alcohol were used as positive controls (137, 138).

### Triton X-100 Subcellular Fractionation

*V. cholerae* cells were subcultured from overnight cultures to an optical density of 0.01 in 50 ml of LB pH 6.5 and grown under Vir Ind for 6-8 hours. Cells were then pelleted by centrifugation (2450 X g 15 minutes), and resuspended in 500 µl of phosphate buffered saline (pH 7.4). Cells were then pelleted by centrifugation (2450 X g 15 minutes).

For spheroplast fractionation, cells were resuspended in 100 µl of 200mM Tris HCl. After resuspension, components were added sequentially to each sample: 200 µl of 200mM Tris HCl and 1M sucrose, 20 µl of 10mM EDTA, 20 µl of lysozyme (10mg/ml), 10 µl of protease inhibitor cocktail (Sigma), and 600 µl of H_2_O. Samples were then incubated at room temperature for 30 minutes. After room temperature incubation 700 µl of 2% Triton X-100, 50mM Tris HCl, and 10mM MgCl_2_ was added.

For gentle cell lysis, pelleted cells were resuspended in 5 ml of Triton X-100 buffer (1% Triton X-100, 10mM imidazole, 500mM HEPES, 10% glycerol, 2M MgCl_2_). Samples then underwent three rounds of freeze-thaw lysis in 180 proof ethanol at -80°C.

Triton X-100 soluble and insoluble membrane fractions were then separated by ultracentrifugation (100,000 X g 1 hour). The supernatant (i.e., the Triton X-100 soluble fraction; TS) and the pellet (i.e., the Triton X-100 insoluble fraction; TI) were collected. The TI fraction was resuspended in 500µl of 2% SDS and 10mM imidazole. The TS fraction was concentrated using Amicon protein concentrators with a 10KDa cutoff (Sigma).

### Co-affinity precipitation

For co-affinity precipitation experiments *V. cholerae* cells were grown as described in the Triton X-100 Subcellular Fractionation section. After cells were suspended in PBS, proteins were cross linked by adding 1mM Dithiobis(succinimidyl propionate) (DPS) to cell suspensions and samples were incubated on ice for 30 minutes. DPS was quenched by adding Tris HCl pH 8.5 to final concentration of 1M and incubating cells on ice for an additional 15 minutes. Cells were then pelleted by centrifugation (2450 X g 15 minutes) and TI and TS fractions were collected via the gentle cell lysis method discussed in the Triton X-100 Subcellular Fractionation section. TI fractions were resuspended in 5ml of 2% sodium dodecyl-sulfate and 10mM imidazole. After collection of TI and TS fractions 100 µl of His-affinity gel (i.e., Ni-NTA Magnetic Agarose Beads) (ZYMO Research) and 10 µl of protease inhibitor cocktail (Sigma) was added to the TI and TS fractions and samples were incubated on a rocking platform overnight at 4°C. TI samples were then incubated at 40°C for 20 minutes to completely solubilize the sample. Samples were then centrifuged (2450 X g 15 minutes) and Ni-NTA agarose beads were washed three times with either Triton X-100 buffer or 2% sodium dodecyl-sulfate and 10mM imidazole. between wash steps TI Ni-NTA agarose beads were incubated at 40°C for 5 minutes. Equal volume of laemmli buffer was added to each sample (BIO-RAD) and then boiled for 5 minutes. Boiled samples were then used directly for western blot analysis.

### Quantification and statistical analysis

Data presented here are arithmetic mean and error bars represent the standard error of the mean, unless otherwise stated. The data were not converted (i.e., log-transformed) prior to statistical analysis. A Shapiro–Wilk test was used to determine if data were normally distributed. Mann-Whitney test was used to measure statistical significance in non-normally distributed data sets. Student’s t-test, one tailed or two tailed, was used to make pairwise comparisons of normally distributed data sets. A P-value ≤0.05 was considered statistically significant. Statistical tests are indicated in each figure legend. Unless otherwise stated, averages represent three or more biological replicates.

## Acknowledgments

This work was supported by the Rudolph Hugh Endowment at Michigan State University (V.J.D.), the Wentworth fellowship, the Rudolph Hugh fellowship, the Michigan State University College of Natural Science Dissertation Continuation and Completion fellowships, and the Integrated Pharmacological Sciences Training Program at MSU (2T32 GM092715), L.M.D. was a trainee. We thank Ron Cook and Dr. Christoph Benning for conducting the FAME analysis of the TI and TS membrane fractions. We thank Dr. Andrew Van Alst, Beth Ottosen, and Rhiannon Leveque for critical review of this manuscript.

