## Supplemental Data for "An essential host dietary fatty acid stimulates TcpH inhibition of TcpP proteolysis enabling virulence gene expression in *Vibrio cholerae*"

*Address correspondence to Victor J. DiRita

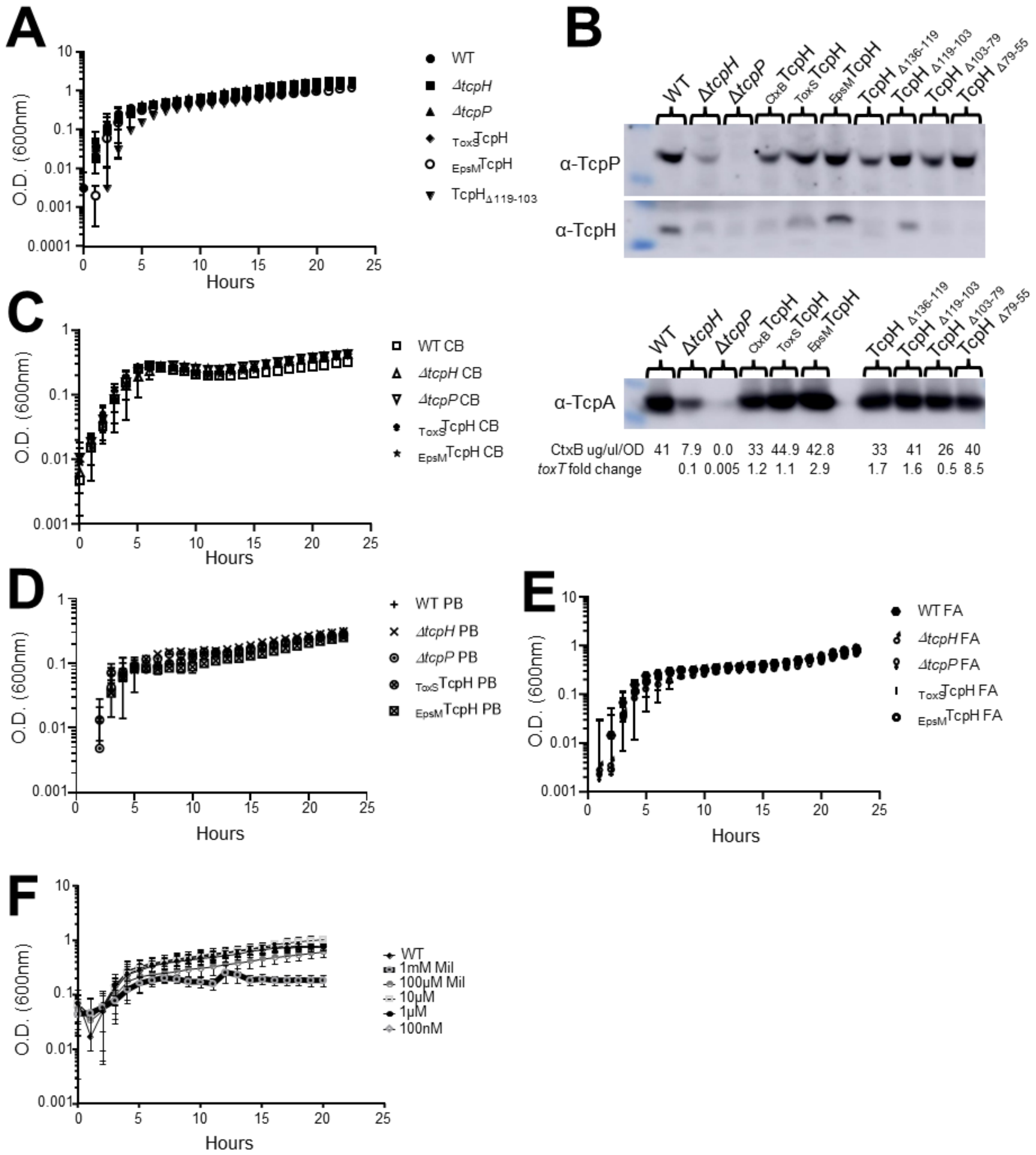


**Figure S1. Growth rates of Transmembrane (TM) and Periplasmic (Peri) TcpH variants are similar to WT cells. Related to Figures 1-3.** A) Virulence inducing conditions growth curve of TcpH TM and Peri constructs respectively. B) *in vitro* characterization of TcpH TM and Peri chromosomal constructs grown under virulence inducing conditions. Western blots of whole-cell lysates probed with α-TcpP (top), α-TcpH (middle), and α-TcpA (bottom). In addition, CtxB levels and *toxT* transcription, relative to WT, were also determined for the TcpH TM and Peri constructs. Average CtxB levels and *toxT* fold change (relative to WT) for each strain are indicated below the western blot. C) Virulence inducing condition growth curve supplemented with crude bile (0.4%). D) Virulence inducing condition growth curve supplemented with purified bile salts (cholate/deoxycholate 100μM). E) Virulence inducing condition growth curve supplemented with linolenic acid (500μM). F) LB, 37°C, growth curve with 1mM to 100nM Miltefosine.

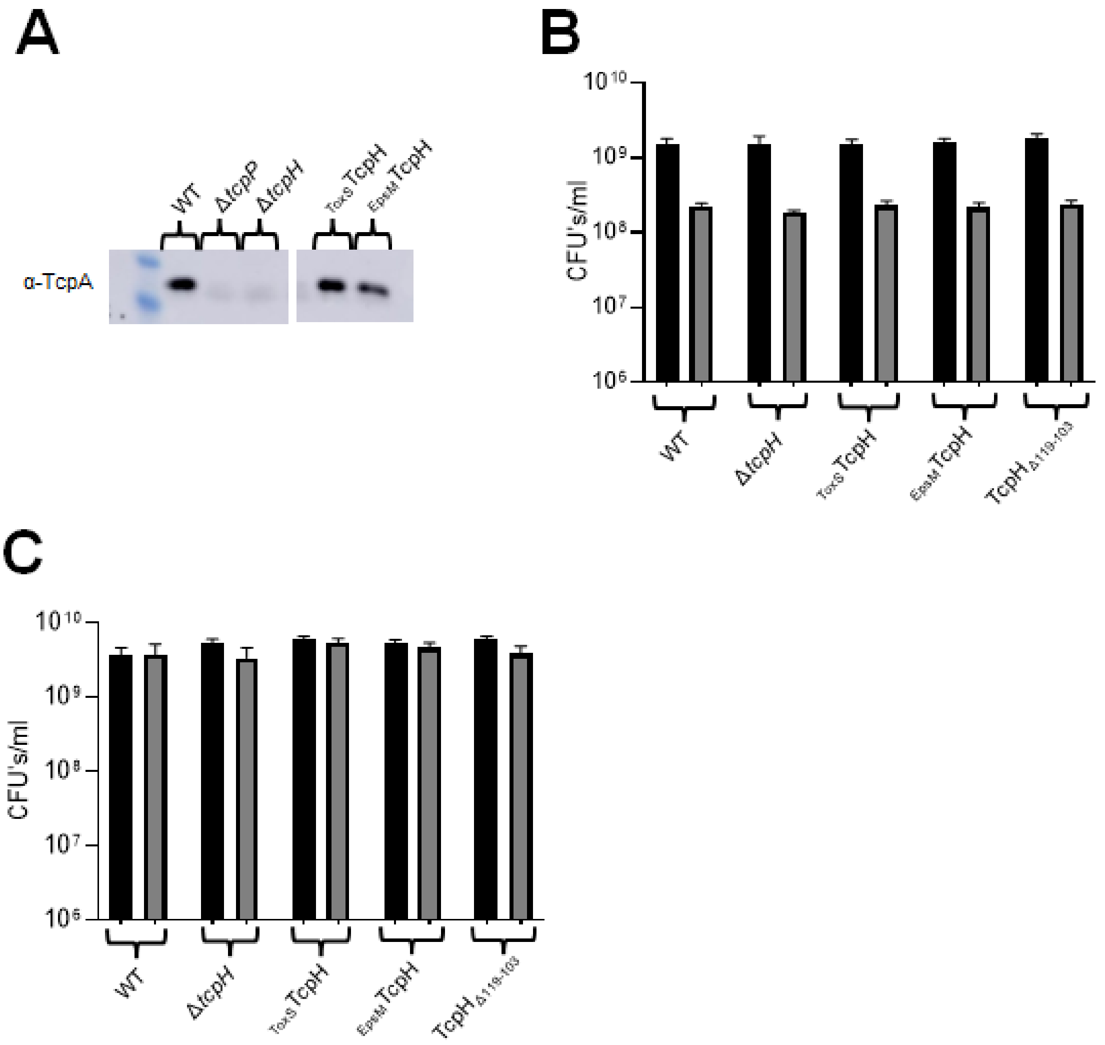


**Figure S2: TcpH transmembrane and periplasmic constructs display WT growth in adult mice feces, and TcpH transmembrane constructs inhibit RIP of TcpP. Related to Figure 2.** A) Western blot of Initial inoculums used to infect infant mice in Figure 2A. B) Filter sterilized mice fecal growth curve. C) Non-filtered (i.e, non-sterile) mice fecal growth curve. Δ*tcpP* was excluded from non-sterile mice fecal growth experiment due to limited supply of non-sterile mice fecal media. Error bars represent standard error of the mean.

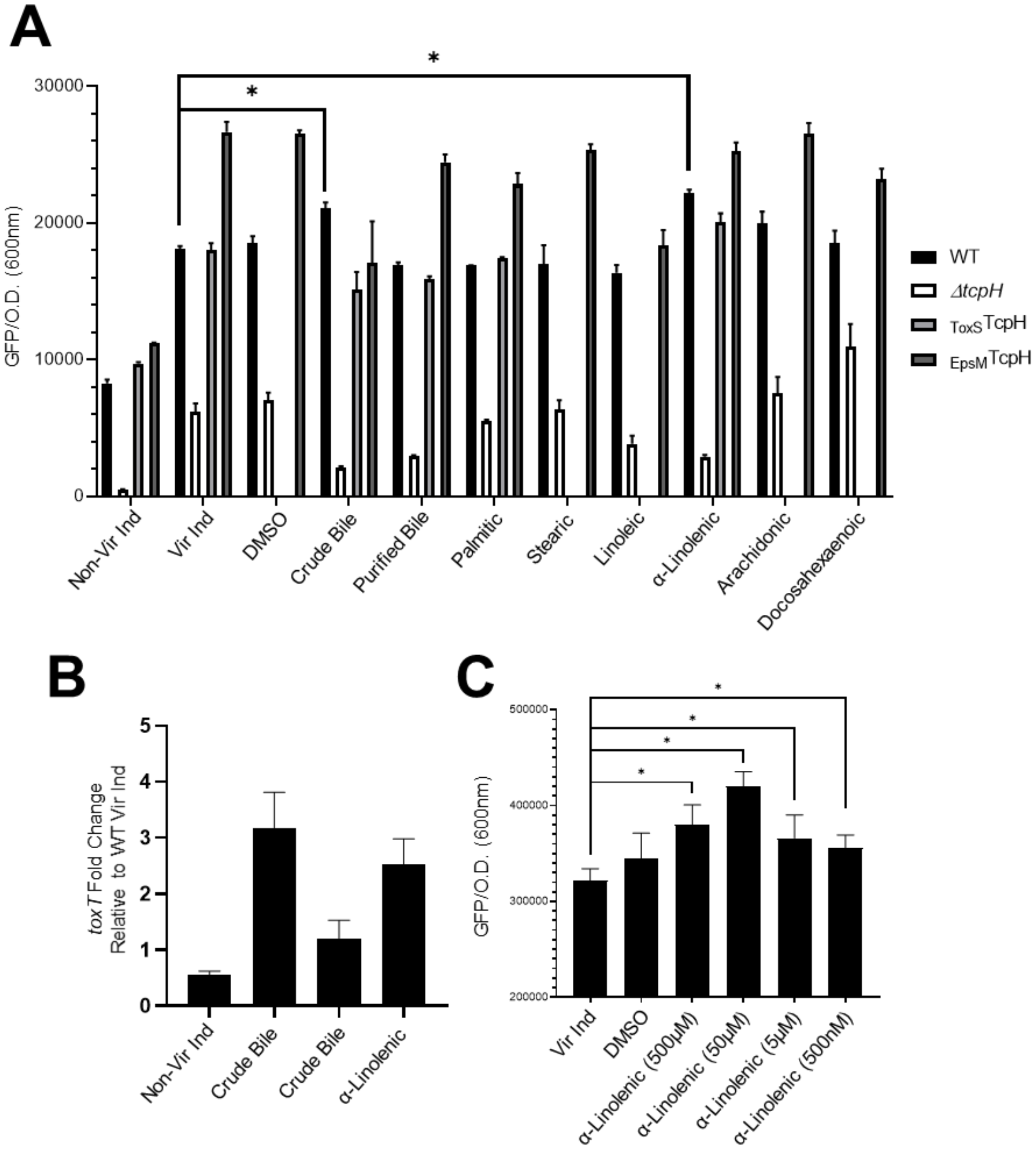


**Figure S3: α-Linolenic acid stimulates *toxT* transcription in a TcpH dependent manner. Related to Figure 3.** A) *toxT* transcription was determined using a plasmid based *toxT::GFP* transcriptional reporter. B) *toxT* transcription in WT *V. cholerae* cells using RT-qPCR, determined via ΔΔCT method. Cells were incubated in Vir Ind for 4hrs and then transferred to indicated conditions for an additional 4hrs. RNA was collected at the 8hr time point. C) *toxT* expression in WT cells determined using a plasmid based *toxT::GFP* transcription reporter. Concentrations of α-linolenic acid (LA) used are displayed below each bar. Lower concentrations of LA (50μM) were tested with control groups (Δ*tcpH* and $_{EpsM}$TcpH) and were not found to increase *toxT* transcription, data not shown. Error bars represent the standard error of the mean. Two-tailed Student's t-test was used to determine statistical significance. *indicates a P-value of < 0.05.

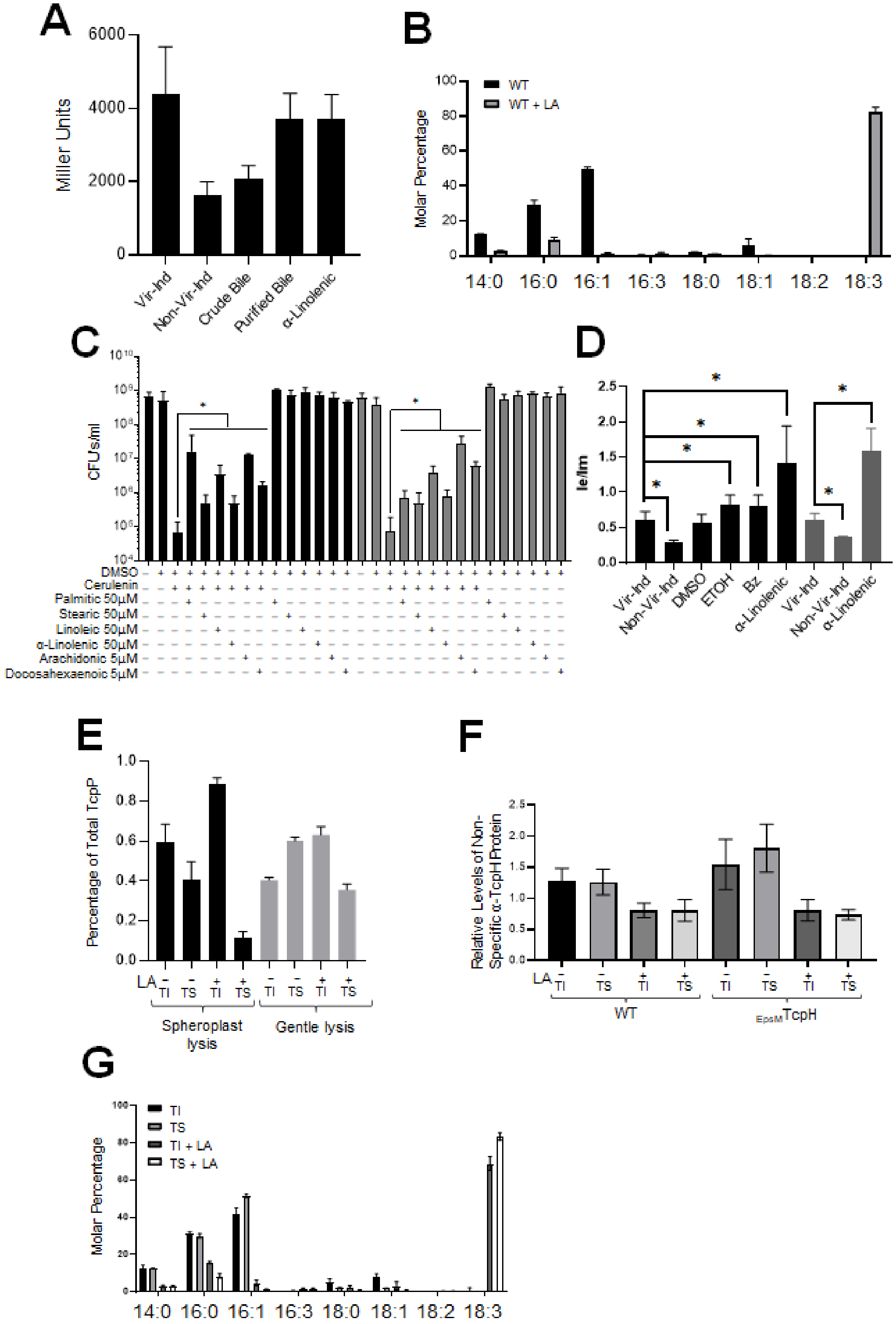


**Figure 4S: α-Linolenic acid is utilized by *V. cholerae* cells, α-linolenic increases membrane fluidity, and does not influence *tcpP* transcription, or promote non-specific protein association within detergent resistant membranes. Related to Figure 3 and 4.** A) *tcpP*

transcription in WT *V. cholerae* cells determined using *tcpP::lacZ* transcription reporter. *tcpP* transcription was determined by quantifying LacZ activity (i.e., calculating Miller Units) (130). B) Molar percentage of fatty acids present in whole *V. cholerae* cells. LA, indicates cells were cultured with α-linolenic acid. Error bars represent the standard deviation, and the average values here represent two biological replicates. C) WT (black bars) and $_{EpsM}$TcpH (grey bars) colony forming units after 24 hours of growth in M9 minimal media (0.05% glucose) at 37°C. The ledged below indicates the presence of additional compounds or fatty acids ("+" indicates present and "-" indicates absence). DMSO and cerulenin final concentration 1% (v/v) and 10 μg/ml respectively. Error bars indicate standard deviation. D) Membrane fluidity of WT (black bars) and $_{EpsM}$TcpH (grey bars) cells grown with and without α-linolenic acid determined by the ratio of excimer (470 nm) and monomer (400 nm) of pyrenedecanoic acid. Error bars represent standard deviation. C and D) a one-tailed Student's t-test was used to determine statistical significance. *indicates a P-value of < 0.05. E) TcpP levels within Triton X-100 soluble (lipid disordered) and Triton X-100 insoluble (lipid ordered) membrane fractions with and without α-linolenic acid supplementation (LA). Black bars indicate TI and TS fractions were collected using the spheroplast method of cell lysis, and gray bars indicate TI and TS factions were collected using the gentle freeze-thaw cell lysis method. ImageJ was used to perform the densitometry analysis. Error bars represent the standard error. F) Relative levels of the non-specific loading control in α-TcpH westerns is equally distributed among Triton soluble (i.e., TS; lipid disordered) and Triton insoluble (i.e., TI; lipid ordered) fractions. Addition of α-linolenic acid (LA, 500μM), indicated by +/-, does not change this distribution. Relative levels of the non-specific loading control were determined via densitometry analysis. Densitometry analysis was conducted using ImageJ. Error bars represent the standard error. G) Molar percentage of fatty acids within WT *V. cholerae* Triton soluble (i.e., TS; lipid disordered) and Triton insoluble (i.e., TI; lipid ordered) fractions. LA, indicates cells were cultured with α-linolenic acid. Error bars represent the standard deviation, and the average values here represent two biological replicates.

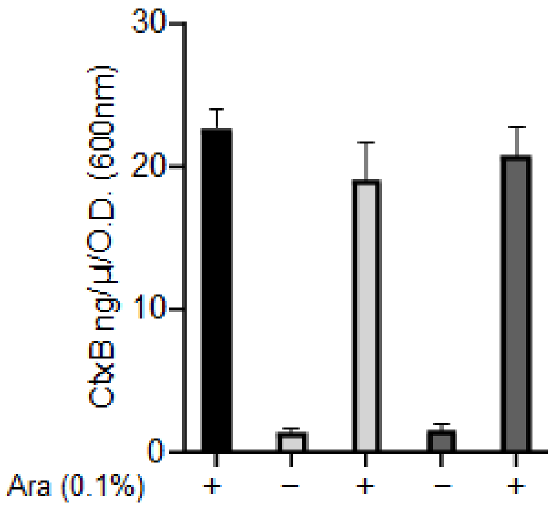


**Figure S5: Hsv-His(6x) tagged TcpP constructs remain functional. Related to Figure 5.** CtxB levels, measured via ELISA, in culture supernatants collected from cultures incubated with *V. cholerae* cells cultured in virulence inducing conditions for 24hrs. Black bars represent WT cells. Light gray bars represent Δ*tcpP* complemented with pBAD18-*Hsv-His(6x)-tcpP* , and dark gray bars represent Δ *tcpP* complemented with pBAD18-*tcpP-His(6x)-Hsv*. *tcpP* constructs were ectopically expressed from pBAD18 using arabinose (Ara 0.1% w/v). + indicates arabinose was added to the culture.

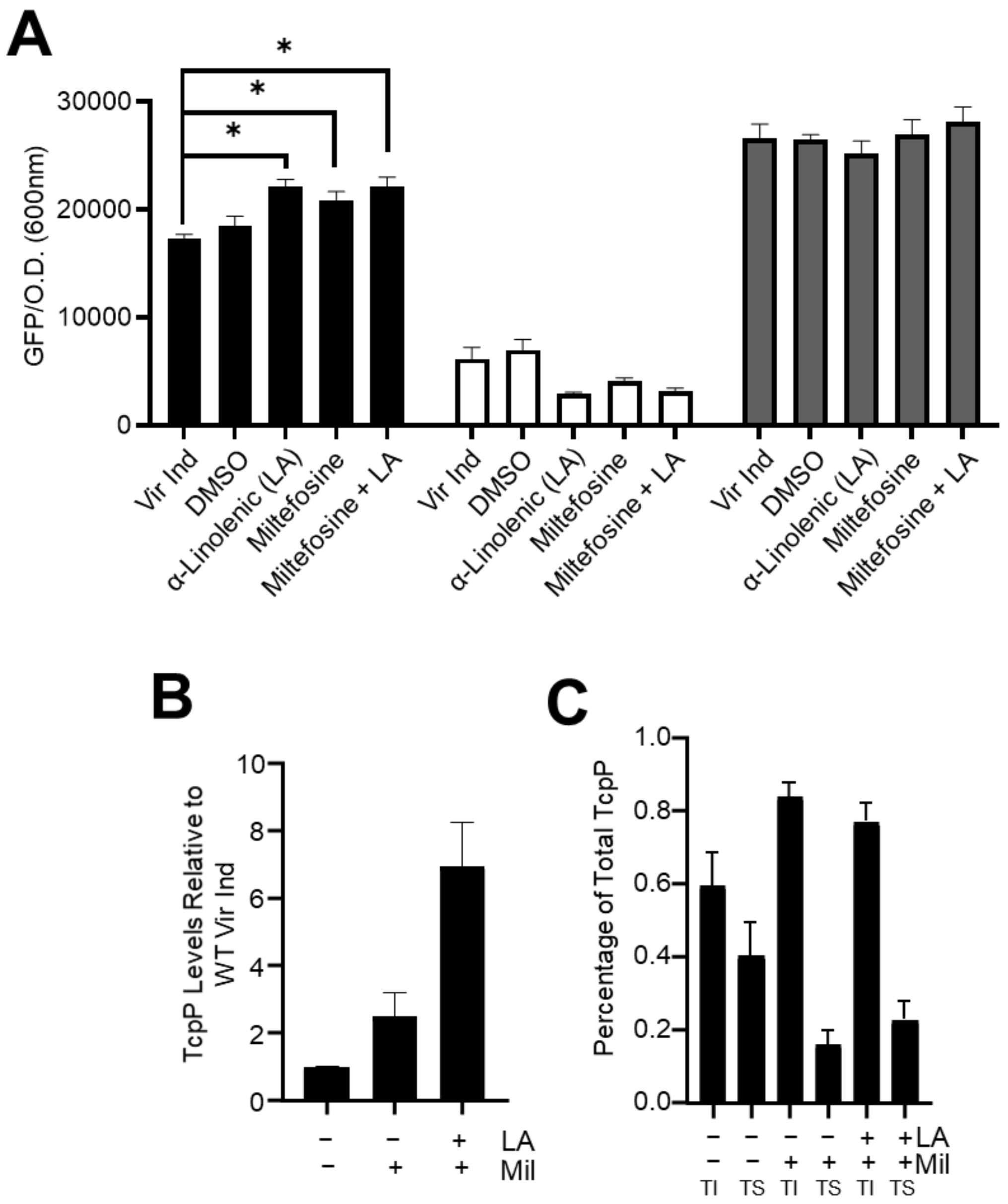


**Figure S6: Miltefosine and α-linolenic acid function synergistically to stimulate *toxT* expression.** A) *toxT* transcription in WT (black bars), Δ*tcpH* (white bars), and $_{EpsM}$TcpH (gray bars) was determined using a plasmid based *toxT::GFP* transcriptional reporter. Error bars represent the standard error of the mean. B) TcpP levels relative to WT *V. cholerae* cells grown in Vir Ind conditions for 8 hours with and without miltefosine and or linolenic acid. C) Percentage of TcpP molecules present in the TI (Triton insoluble; lipid ordered membrane domain) and TS (Triton soluble; lipid disordered membrane domain) membrane fractions within WT *V. cholerae* cells. B and C) Densitometry analysis was done using ImageJ to quantify TcpP levels. Error bars represent standard error of the mean. LA: α-linolenic acid (500µM) Mil: miltefosine (10µM). A two-tailed Student's t-test was used to determine statistical significance. *indicates a P-value of < 0.05.

**Table S1:** strains used in this study.

| Strain | Description | Reference |
|---|---|---|
| *V. cholerae* 0395 classical biotype | Wild type | DiRita lab collection |
| *V. cholerae* Δ*tcpH* | Isogenic deletion | DiRita lab collection |
| *V. cholerae* Δ*tcpP* | Isogenic deletion | DiRita lab collection |
| *V. cholerae* Δ*tcpH* pBAD18-empty vector | Overexpression plasmid vector | DiRita lab collection |
| *V. cholerae* Δ*tcpH* pBAD18 TcpH | Δ*tcpH* complementation with ectopic *tcpH* | This study |
| *V. cholerae* Δ*tcpPH* pBAD18 $_{CtxB}$TcpH | Δ*tcpH* complementation with ectopic *tcpH* TM construct | This study |
| *V. cholerae* Δ*tcpH*; Δ*yaeL* pBAD18-empty vector | Overexpression plasmid | This study |
| *V. cholerae* Δ*tcpH*; Δ*yaeL* pBAD18 $_{CtxB}$TcpH | Δ*tcpH* complementation with ectopic *tcpH* TM construct | This study |
| *V. cholerae* Δ*tcpH*; Δ*yaeL* pBAD18 $_{ToxS}$TcpH | Δ*tcpH* complementation with ectopic *tcpH* TM construct | This study |
| *V. cholerae* Δ*tcpH*; Δ*yaeL* pBAD18 $_{EpsM}$TcpH | Δ*tcpH* complementation with ectopic *tcpH* TM construct | This study |
| *V. cholerae* Δ*tcpH*; Δ*yaeL* pBAD18 $TcpH_{\Delta 136-119}$ | Δ*tcpH* complementation with ectopic *tcpH* Peri construct | This study |

| | | |
|---|---|---|
| *V. cholerae* Δ*tcpH*; Δ*yaeL* pBAD18 $TcpH_{\Delta 136\text{-}103}$ | Δ*tcpH* complementation with ectopic *tcpH* Peri construct | This study |
| *V. cholerae* Δ*tcpP* pBAD18 *Hsv-His(6x)-tcpP* | N-terminal *tcpP* co-immuno precipitation construct | This study |
| *V. cholerae* Δ*tcpP* pBAD18 *tcpP-His(6x)-Hsv* | C-terminal *tcpP* co-immuno precipitation construct | This study |
| *V. cholerae* Δ*tcpH* pBAD18 *Hsv-His(6x)-tcpH* | N-terminal *tcpH* co-immuno precipitation construct | This study |
| *V. cholerae* Δ*tcpH* pBAD18 *tcpH-His(6x)-Hsv* | C-terminal *tcpH* co-immuno precipitation construct | This study |
| *V. cholerae* Δ*yaeL* pBAD18 *Hsv-His(6x)-tcpP* | N-terminal *tcpP* co-immuno precipitation construct | This study |
| *V. cholerae* Δ*yaeL* pBAD18 *tcpP-His(6x)-Hsv* | C-terminal *tcpP* co-immuno precipitation construct | This study |
| *V. cholerae* $_{CtxB}$TcpH | chromosomal construct | This study |
| *V. cholerae* $_{ToxS}$TcpH | chromosomal construct | This study |
| *V. cholerae* $_{EpsM}$TcpH | chromosomal construct | This study |
| *V. cholerae* $TcpH_{\Delta 136\text{-}119}$ | chromosomal construct | This study |
| *V. cholerae* $TcpH_{\Delta 136\text{-}103}$ | chromosomal construct | This study |
| *V. cholerae* $TcpH_{\Delta 119\text{-}103}$ | chromosomal construct | This study |
| *V. cholerae* $TcpH_{\Delta 103\text{-}79}$ | chromosomal construct | This study |

| | | |
|---|---|---|
| *V. cholerae* $TcpH_{\Delta 79\text{-}55}$ | chromosomal construct | This study |
| *V. cholerae* TcpHC114S | isogenic mutant | This study |
| *V. cholerae* TcpHC114S/C132S | isogenic mutant | This study |
| *V. cholerae* pBH6119-*toxT::GFP* | *toxT* transcription reporter | Anthouard R, and DiRita VJ. mBio. 2013. |
| *V. cholerae* Δ*tcpH* pBH6119-*toxT::GFP* | *toxT* transcription reporter | This study |
| *V. cholerae* Δ*tcpP* pBH6119-*toxT::GFP* | *toxT* transcription reporter | This study |
| *V. cholerae* $_{CtxB}$TcpH pBH6119-*toxT::GFP* | *toxT* transcription reporter | This study |
| *V. cholerae* $_{ToxS}$TcpH pBH6119-*toxT::GFP* | *toxT* transcription reporter | This study |
| *V. cholerae* $_{EpsM}$TcpH pBH6119-*toxT::GFP* | *toxT* transcription reporter | This study |
| *V. cholerae* $TcpH_{\Delta 136\text{-}119}$ pBH6119-*toxT::GFP* | *toxT* transcription reporter | This study |
| *V. cholerae* $TcpH_{\Delta 136\text{-}103}$ pBH6119-*toxT::GFP* | *toxT* transcription reporter | This study |
| *V. cholerae* $TcpH_{\Delta 119\text{-}103}$ pBH6119-*toxT::GFP* | *toxT* transcription reporter | This study |

| | | |
|---|---|---|
| *V. cholerae* TcpH$_{\Delta 103\text{-}79}$<br>pBH6119-*toxT::GFP* | *toxT* transcription reporter | This study |
| *V. cholerae* TcpH$_{\Delta 79\text{-}55}$<br>pBH6119-*toxT::GFP* | *toxT* transcription reporter | This study |
| *V. cholerae* TcpHC114S<br>pBH6119-*toxT::GFP* | *toxT* transcription reporter | This study |
| *V. cholerae* TcpHC114S/C132S<br>pBH6119-*toxT::GFP* | *toxT* transcription reporter | This study |
| *E. coli* ET12567 Δ*dapA* | Cloning vector recipient | Allard, N., et. al. 2015. Canadian Journal of Microbiology, 61(8), pp.565-574. |
| *E. coli* ET12567 Δ*dapA*<br>pKAS32-empty vector | Plasmid vector strain | DiRita lab collection |
| *E. coli* ET12567 Δ*dapA*<br>pBAD18-empty vector | Plasmid vector strain | DiRita lab collection |

**Table S2:** All primers used in this study contain Kpn1-HiFi (forward primers) and Xba1 (reverse primers) restriction sites.

| Description | Sequence |
|---|---|
| pKAS FW | gcctctaaggttttaagt |
| pKAS RV | ctttcaaggtagcggttacc |
| pBAD18 FW | ctgtttctccatacccgtt |
| pBAD18 RV | ggctgaaaatcttctct |
| pKAS-TcpP promoter FW | ctaacgttaacaaccggtactttcgagtgatagaaaaagg |
| pKAS-TcpP FW | ctaacgttaacaaccggtacatggggtatgtccgcgtg |
| pKAS-downstream TcpH RV | aaatttgcgcatgctagctatagttcttggtctttttagataacgtaagc |
| TcpP-CtxBss FW | atgcactaaaaattaaaagacattagaatgattaaattaaaatttgg |
| TcpP-CtxBss RV | aatttaatcattctaatgtcttttaatttttagtgcattctaatgtcttc |
| CtxBss-TcpHperi FW | tcttcagcatatgcacatggaccgatgcgacaaaaaaac |
| CtxBss-TcpHperi RV | gtcgcatcggtccatgtgcatatgctgaaga |
| TcpP-EpsMss FW | atgcactaaaaattaaaagacattagaatgatgaaagaattattggctc |
| TcpP-EpsMss RV | tctaatgtcttttaatttttagtgcattctaatgtcttc |
| EpsMss-TcpHperi FW | gggaatatggccgatgcgacaaaaaaac |
| EpsMss-TcpHperi RV | gtcgcatcggccatattccccaataagc |

| | |
|---|---|
| TcpP-ToxSss FW | atgcactaaaaattaaaagacattagaatgcaaaatagacacatcg |
| TcpP-ToxSss RV | cgatgtgtctattttgcattctaatgtcttttaatttttagtgcattctaatgtcttc |
| ToxSss-TcpHperi FW | ttgggggagtccgatgcgacaaaaaaac |
| ToxSss-TcpHperi RV | tgcatgcctgcaggtcgactctaaaaatcgctttgacag |
| $TcpH_{\Delta 136\text{-}119}$ FW | cgccttcccttagggtcttatcatgagccgc |
| $TcpH_{\Delta 136\text{-}119}$ RV | tgataagaccctaagggaaggcgagaaaacaac |
| $TcpH_{\Delta 136\text{-}103}$ FW | tgattacaattagggtcttatcatgagccgc |
| $TcpH_{\Delta 136\text{-}103}$ RV | tgataagaccctaattgtaatcacggctcacattactttc |
| $TcpH_{\Delta 119\text{-}103}$ FW | tgattacaattacaagcagcttacggctg |
| $TcpH_{\Delta 119\text{-}103}$ RV | taagctgcttgtaattgtaatcacggctcac |
| $TcpH_{\Delta 103\text{-}79}$ FW | tcaaacattggtgttgagtatttatcaactc |
| $TcpH_{\Delta 103\text{-}79}$ RV | tactcaacaccaatgtttgataacgtgtag |
| $TcpH_{\Delta 79\text{-}55}$ FW | taatctatccccagatcctagctctcag |
| $TcpH_{\Delta 79\text{-}55}$ RV | taggatctggggatagattaccttgataagtag |
| TcpHC114S FW | tcaactcggcaaaggtagttttctcgccttccc |
| TcpHC114S RV | gggaaggcgagaaaactacctttgccgagttga |
| TcpHC132S FW | ggttttccagtcaaagcgatttttag |
| TcpHC132S RV | ctaaaaatcgctttgactggaaaacc |

| | |
|---|---|
| pBAD18-CtxBss FW | agcgaattcgagctcggtaccaaagggagcattataagacattagaatgattaaattaaaatttgg |
| pBAD18-ToxSss RV | agcgaattcgagctcggtaccaaagggagcattatatgcaaaatagacacatcg |
| pBAD18-EpsMss FW | agcgaattcgagctcggtaccaaagggagcattatatgatgaaagaattattggctc |
| pBAD18-TcpH FW | agcgaattcgagctcggtaccaaagggagcattatatgcacaaaaaattaaaagcttg |
| pBAD18-TcpH RV | tgcatgcctgcaggtcgactctaaaaatcgctttgacag |
| pBAD18-TcpH$_{\Delta 136\text{-}119}$ RV | tgcatgcctgcaggtcgactctaagggaaggcgagaaaacaac |
| pBAD18-TcpH$_{\Delta 136\text{-}103}$ RV | tgcatgcctgcaggtcgactctaattgtaatcacggctcacattactttc |
| pBAD18 Hsv-His(6x) FW | ttcgagctcggtaccaaagggagcattatatgcagccggaactggcgccggaagatccg |
| Hsv-His(6x)-TcpP FW | ccggaagatccggaagattgccatcatcatcatcatcatatggggtatgtccgcgtg |
| Hsv-His(6x)-TcpP RV | cagttccggctgatgatgatgatgatgatgattttttgtgcattctaatgtcttc |
| pBAD18-TcpP RV | tgcatgcctgcaggtcgactttaatttttgtgcattctaatgtcttctgttc |
| pKT25-TcpP FW | ggctgcagggtcgactatggggtatgtccgc |
| pKT25-TcpP RV | attcttacttacttaggtacttaatttttgtgcattctaatgtcttctgttc |
| pUT18C-TcpH FW | aacgccactgcaggtcgactcagcggtggtggaggttcgaaatgcacaaaaaattaaaag |
| pUT18C-TcpH RV | gatgaattcgagctcggtacctaaaaatcgctttgacaggaaaacc |
| *recA* FW RT-qPCR | attgaaggcgaaatgggcgatag |
| *recA* RV RT-qPCR | tacacatacagttggattgcttg agg |

| | |
|---|---|
| *toxT* FW RT-qPCR | actgatgatcttgatgctatggag |
| *toxT* RV RT-qPCR | catccgattcgttcttaattcacc |
| *tcpP* FW RT-qPCR | tgagtgggggaagataaacg |
| *tcpP* RV RT-qPCR | ttggattgttatccccggta |
